# *In vitro* reconstitution reveals membrane clustering and RNA recruitment by the enteroviral AAA+ ATPase 2C

**DOI:** 10.1101/2023.10.11.561830

**Authors:** Kasturika Shankar, Marie N. Sorin, Himanshu Sharma, Oskar Skoglund, Selma Dahmane, Josy Ter Beek, Solomon Tesfalidet, Louise Nenzén, Lars-Anders Carlson

## Abstract

Enteroviruses are a vast genus of positive-sense RNA viruses that cause diseases ranging from common cold to poliomyelitis and viral myocarditis. They encode a membrane-bound AAA+ ATPase, 2C, that has been suggested to serve several roles in virus replication, e.g. as an RNA helicase and capsid assembly factor. Here, we report the reconstitution of full-length, poliovirus 2C’s association with membranes. We show that the N-terminal membrane-binding domain of 2C contains a conserved glycine, which is suggested by structure predictions to divides the domain into two amphipathic helix regions, which we name AH1 and AH2. AH2 is the main mediator of 2C oligomerization, and is necessary and sufficient for its membrane binding. AH1 is the main mediator of a novel function of 2C: clustering of membranes. Cryo-electron tomography reveal that several 2C copies mediate this function by localizing to vesicle-vesicle interfaces. 2C-mediated clustering is partially outcompeted by RNA, suggesting a way by which 2C can switch from an early role in coalescing replication organelles and lipid droplets, to a later role where 2C assists RNA replication and particle assembly. 2C is sufficient to recruit RNA to membranes, with a preference for double-stranded RNA (the replicating form of the viral genome). Finally, the *in vitro* reconstitution revealed that full-length, membrane-bound 2C has ATPase activity and ATP-independent, single-strand ribonuclease activity, but no detectable helicase activity. Together, this study suggests novel roles for 2C in membrane clustering, RNA membrane recruitment and cleavage, and calls into question a role of 2C as an RNA helicase. The reconstitution of functional, 2C-decorated vesicles provides a platform for further biochemical studies into this protein and its roles in enterovirus replication.

## Introduction

The *Enterovirus* genus (family: *Picornaviridae*) are positive-sense single-stranded RNA (+ssRNA) viruses, whose ∼7.5kb genome is encapsidated in a 30 nm non-enveloped, icosahedral capsid [1, 2]. There are hundreds of known human enteroviruses, responsible for diseases like poliomyelitis (poliovirus, PV), viral myocarditis (Coxsackievirus B3, CVB3), common cold (Rhinoviruses, RVs) and hand, foot and mouth disease (e.g., Coxsackievirus A16 and Enterovirus 71, EV71) [3–5]. In addition, their high mutation rate and frequent recombinations mean that novel enteroviruses emerge regularly [6–8]. PV, which causes the paralytic disease poliomyelitis, has been a major focus of enterovirus research and worldwide vaccination efforts. Since the launch of the Global Polio Eradication Initiative in 1988, the number of poliomyelitis cases have reduced to near-zero in most of the world, but occasional cases of vaccine-derived poliomyelitis highlight the difficulty in achieving complete eradication[9–11]. There is limited progress in the production of vaccines against the many non-polio enteroviruses that cause frequent outbreaks in different parts of the world [12]. Development of pan-enteroviral drugs that inhibit conserved steps of the replication cycle is thus of importance not only for the last phase of polio eradication, but also for minimizing the risk associated with future enterovirus outbreaks, as well as for treating chronic enterovirus infections [13–16].

Enterovirus genome replication is a highly conserved process requiring the formation of cytoplasmic replication organelles (ROs) [17–19]. The ROs are formed from extensively modified cytoplasmic membranes which are turned into an assembly line for viral RNA and virus particles [20, 21]. RO membranes are obtained from several cellular sources. Initially, they come from the disassembly of the Golgi apparatus as well as from utilization of cellular cholesterol [22, 23]. In addition, ROs are enriched by fatty acids from lipid droplets, through RO-lipid droplets contacts mediated by the viral protein 2C [24]. At later time points, induction of macro-autophagy (henceforth: autophagy) provides another source of RO membranes in the form of double-membrane vesicles [25–27]. The direct dependence of enterovirus RNA replication on membranes is supported e.g. by electron microscopy autoradiography, showing that newly synthesized RNA co-localizes with RO membranes [28]. Further, subcellular fractions active in viral RNA synthesis contain membranes, and their activity is inhibited if the membranes are disintegrated by detergents [29, 30]. How viral RNA, viral proteins, and host factors arrange spatially on the cytoplasmic face of RO membranes is not known.

The enterovirus genome has one major open reading frame which is translated as a single major polyprotein, with some enteroviruses also having a smaller accessory open reading frame [31]. The major polyprotein is subsequently cleaved into eleven individual proteins by the viral proteases 2A and 3C (Fig. 1A). Additionally, some important and distinct functions are served by relatively stable, incompletely cleaved products such as 2BC [32]. The non-structural proteins, contained in polyprotein regions P2 and P3, encode functions necessary for RO formation and genome replication (Fig. 1A). Amongst these is the protein 2C, a conserved, multifunctional ATPase of the AAA+ superfamily [33–35]. 2C is essential for both RNA replication and virion assembly, but its precise roles in these processes are unknown [36–40]. 2C and 2BC are sufficient for intracellular membrane rearrangements similar to those observed in infection[41], and 2BC in combination with 3A induces an autophagy response similar to that seen in infection [42]. 2C has a central ATPase domain followed by a Zn finger and a C-terminal α-helix which engages a hydrophobic pocket of another 2C monomer to aid its oligomerization [43–46] (Fig. 1A). ATP hydrolysis by 2C is stimulated by its oligomerization [45, 47], and by binding to single-stranded RNA [48]. Interestingly, even though homologous proteins from other viruses are helicases, many publications report that 2C only has ATP-independent RNA chaperoning activity, and little-to-no ATP-dependent RNA helicase activity, with one study suggesting that post-translational modifications may activate 2C as a helicase [48, 49]. However, it must be noted that biochemical studies on 2C have to date been carried out in simplified systems, using e.g. truncated 2C constructs, fusion proteins, and/or omitting the membrane platform on which 2C is located during infection.

**Figure 1:**
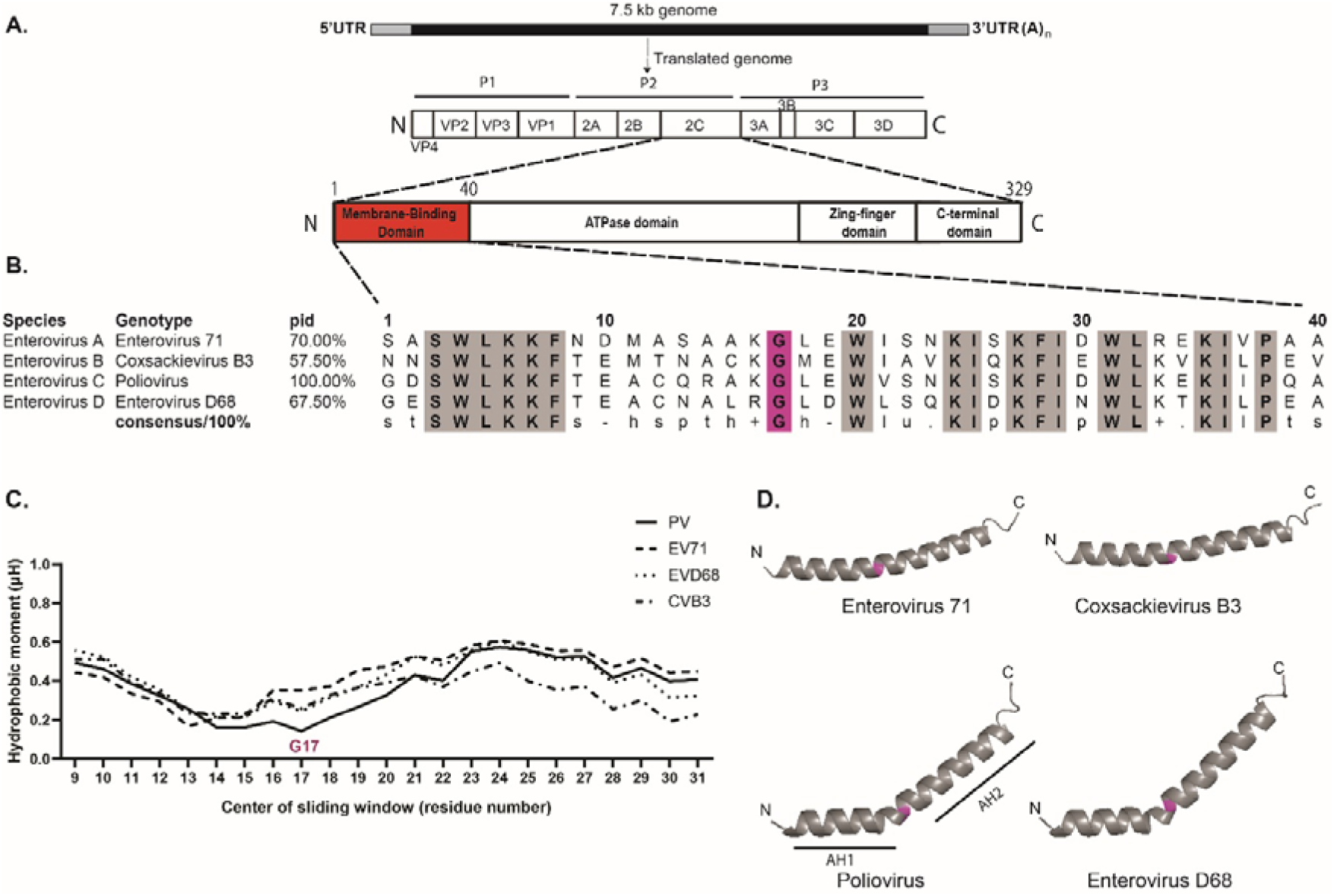
The membrane-binding region of 2C consists of two amphipathic helix regions, separated by a conserved glycine. **(A)** Schematic representation of the *Enterovirus* genome, the translated polyprotein with individual proteins marked, and the 2C protein with known domains marked. **(B)** Multiple sequence alignment of the membrane-binding domain (MBD, residues 1-40) of 2C from representative genotypes of different *Enterovirus* species. Conserved residues are highlighted in grey (s: small residue, t: turn like residue, h: hydrophobic residue, p: polar residue, l: aliphatic residue, +: positively charged residue, -: negatively charged residue). The conserved glycine (number 17 in poliovirus 2C) is highlighted in pink. **(C)** Hydrophobic moment (µH), a measure of the local amphiphilicity of a helix in a 18-residue sliding window, is plotted for representative genotypes of different enterovirus species. The higher the µH value, the more an amphipathic the helix at that position. The position of the conserved glycine 17 is marked. **(D)** Alphafold2 structure prediction of the MBD of 2C from representative genotypes of different enterovirus species. The conserved glycine 17 is highlighted in pink separating AH1 and AH2.

The conserved, N-terminal membrane-binding domain (MBD) of 2C is necessary for virus replication [50]. It has been suggested to contain an amphipathic helix, and has been shown to be necessary and sufficient for membrane association when overexpressed in cells [50–52]. The MBD has further been described to mediate the oligomerization of poliovirus 2C [47], while the C-terminal α-helix appears sufficient to mediate oligomerization of 2C from some other enteroviruses [44]. Due to the hydrophobic nature of the MBD, full-length 2C tends to aggregate during purification, thus prohibiting its use in biochemical experiments. This problem has typically been circumvented by working with a truncated 2C construct lacking the MBD [43–45], or by using high concentrations of detergents [48]. Alternatively, fusing the N-terminus of full-length 2C to the large, hydrophilic maltose-binding protein (MBP) has also been reported to result in a soluble protein [47]. While all of these studies have uncovered important aspects of 2C’s function, they were not able to address the mechanisms of its membrane binding, nor its function as a full-length protein on a membrane.

Here, we biochemically reconstituted the binding of purified, full-length poliovirus 2C to synthetic vesicles. We suggest that the MBD is in fact composed of two amphipathic helix segments, AH1 and AH2, interrupted by an absolutely conserved glycine residue. Both AH1 and AH2 are necessary for complete hexamerization of 2C, whereas AH2 is sufficient for its membrane localization. A novel function for AH1 is identified in tethering membranes together, and we show that 2C’s membrane tethering can be partially outcompeted by nucleic acid. We show that 2C is sufficient to recruit and retain RNA to membranes, and that it has a strong preference for retaining double-stranded RNA at a membrane. Together, these findings provide a biochemical basis for several roles of 2C in enterovirus replication, and establishes a more physiologically relevant biochemical system as a basis for further investigations.

## Results

### The oligomerization state of 2C is controlled by its N-terminal, bipartite amphipathic helix

The membrane-binding domain (MBD) of poliovirus 2C (Fig. 1A) consists of residues ∼1-40 and has been characterized as an amphipathic region [50–54], but details of its organization are not known. We took a computational approach to studying the MBD across enteroviruses. A multiple sequence alignment revealed that several hydrophobic residues (W4, L5, F8, G17, W20, I25, F28, I29, W31, L32 and I36), polar residues (S3, K6, K7, K24, K27, and K35) and P38 are broadly conserved between enterovirus species (Fig. 1B). We next studied the amphipathic nature of the 1-40 region by calculating its hydrophobic moment (µH) in a sliding window of 18 residues. A high value for hydrophobic moment means that a helix is amphipathic (i.e. that one side is more hydrophobic than the other). The calculation suggested that the MBD may, in fact, consist of two separate amphipathic helices, with the space between the amphipathic regions coinciding with the conserved residue G17 (Fig. 1C). The amphipathic nature of these two regions was corroborated by a helical wheel plot (Fig.S1A). Since glycine is a helix breaker residue, we hypothesized that G17 in fact divides the MBD into two amphipathic helices, AH1 and AH2. AlphaFold2 [55] structure predictions of 2C(1-40) of several enteroviruses indeed suggested that a kink at G17 is a conserved structural feature (Fig. 1D). The extent of the kink around G17 may not be accurately predicted for a single 2C sequence, as this feature may in fact be flexible, but the structure predictions more robustly indicated that AH1 has a hydrophobic surface which is offset from AH2 [56] (Fig. S1B). Together, these data suggest that the MBD of 2C consists of two amphipathic helix regions, AH1 and AH2, and we thus set out to study their individual roles in the protein.

Recombinant full-length 2C has been reported to be insoluble in the absence of detergents unless its N-terminus is fused to a solubility-enhancing tag [43, 48]. We thus purified the N-terminally His_6_MBP-tagged, full-length poliovirus 2C protein (residues 1-329) and its N-terminally truncated variants 2C(ΔAH1) (residues 12-329), 2C(ΔAH2) (residues 1-11, 41-329), 2C(ΔMBD) (residues 41-329) and 2C(Δ115) (residues 116-329). The constructs 2C(ΔAH1), 2C(ΔAH2) and 2C(ΔMBD) correspond to the truncation of AH1, AH2, and AH1-2, respectively, whereas 2C(Δ115) corresponds to a soluble fragment which has previously been crystallized (Fig. 2A, B, Fig. S2) [43]. Full-length 2C has been suggested to be a homo-oligomer, possibly a hexamer [43, 45, 47, 57, 58]. We wanted to study the individual roles of the AH1 and AH2 regions in 2C oligomerization. Notably, every N-terminal truncation led to a gradual shift towards higher elution volumes in size exclusion chromatography (Fig. 2B-C), indicating that every part of the MBD plays a role in 2C oligomerization. To further dissect the role of AH1 and AH2, we determined the molecular mass of full-length and truncated 2C constructs. Mass photometry of MBP-2C at 100 nM revealed presence of monomeric protein, but also high molecular-mass species at 469±180 kDa (Fig. 2B,D,S3). The peak value of 469 kDa correlated well with the theoretical mass of an MBP-2C hexamer at 484.2 kDa. Since the MBP-2C peak appeared broad, we compared it to apoferritin, a rigid, 474 kDa homo-oligomeric protein complex. Mass photometry of apoferritin at the same concentration (100 nM) indeed resulted in narrower, single distribution of 460±62 kDa, suggesting that the oligomeric form of MBP-2C is predominantly, but not solely, a hexamer under the conditions of the assay (Fig. 2D, Fig. S3A-B). Mass photometry of MBP-2C(ΔAH1) revealed a slightly shifted oligomer peak at 375±246 kDa, which may correspond to a more heterogeneous oligomeric state with a higher proportion of small species (Fig. 2B,E, Fig. S3C). MBP-2C(ΔAH2) existed as a mix of monomer and dimer (Fig.2B,F, Fig. S3D). The shorter constructs 2C(ΔMBD) and 2C(Δ115) were stable in solution even after removal of the MBP tag, and their masses were analyzed at solution concentrations of 38 µM and 134 µM, respectively, by size exclusion chromatography coupled to multi-angle light scattering (SEC-MALS). The mass estimates throughout the predominant peak for two consecutive runs were 43 kDa and 52 kDa for 2C(ΔMBD), i.e. intermediate between monomer (at 32.8 kDa) and dimer (Fig. 2G), possibly due to an equilibrium between the two states. The predominant peak for two SEC-MALS runs for 2C(Δ115) gave an estimated mass of 27 kDa and 22 kDa, consistent with the expected 23.8 kDa of a monomer (Fig. 2H). The MBP-fused versions of these constructs both had SEC-MALS mass estimates corresponding to monomers (Fig. S4A-B).

**Figure 2:**
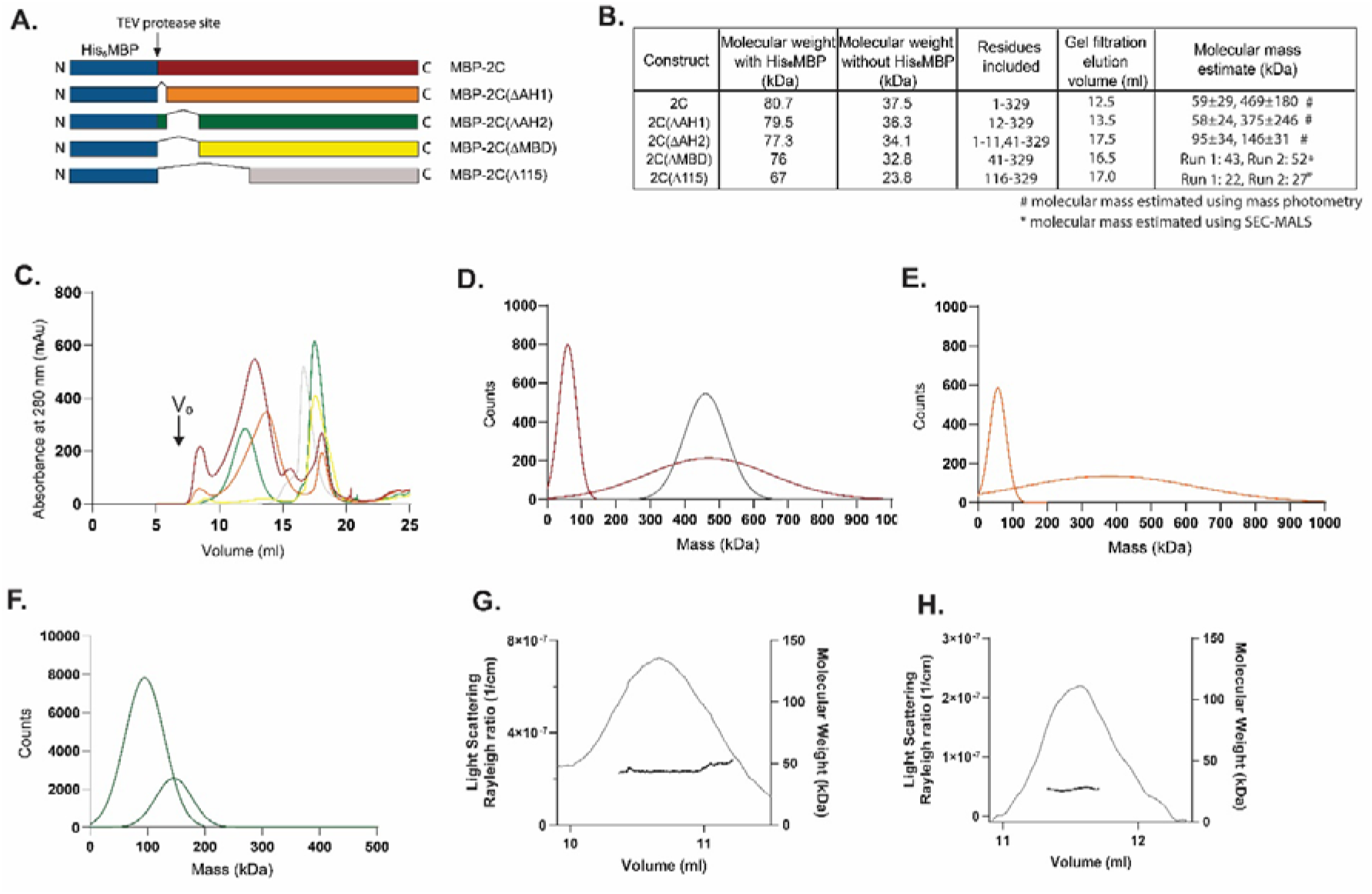
The membrane-binding domain regulates 2C oligomerization. **(A)** Schematic representation of all His_6_-MBP-(TEV) constructs used in the paper: full-length 2C, and N-terminally truncated versions: 2C(ΔAH1) corresponding to deletion of AH1, 2C(ΔAH2) corresponding to deletion of AH2, 2C(ΔMBD) corresponding to deletion of AH1-2 (entire MBD), and 2C(Δ115) corresponding to deletion of MBD and part of the ATPase domain (comparable to published crystal structure)[43]. **(B)** Table showing monomer molecular mass of different constructs with and without MBP, and the 2C residues included in the construct. The table also lists the elution volumes in the size exclusion chromatography in (C) and the mass estimates from mass photometry and SEL-MALS as indicated. **(C)** Elution profiles of the fluorophore-labeled constructs described in panel A on a Superose 6 Increase size-exclusion column. Colors as in (A). The peaks that were used for subsequent analyses were centered at the elution volumes listed in (B). The peak for MBP-2C(ΔAH2) at 12.0 ml only appeared upon fluorophore-labeling and was not used in the biochemical assay. V_0_, column void volume. **(D-F)** Molecular mass distribution (Gaussian fits to mass photometry raw data shown in Fig. S3) at 100 nM concentration of MBP-2C and apoferritin (apoferritin black) (D), MBP-2C(ΔAH1) (E) and MBP-2C(ΔAH2) (F). **(G-H)** SEC-MALS data showing estimated molecular mass of MBP-free 2C(ΔMBD) and 2C(Δ115) at concentrations of 38 and 134 μM, respectively. Throughout the main elution peak, 2C(ΔMBD) had an estimated mass of 43 and 52 kDa for two consecutive runs, whereas 2C(Δ115) had an estimated mass of 22 and 27 kDa for two consecutive runs.

Taken together, structure predictions indicate that the conserved glycine 17 divides the MBD into two amphipathic helices, AH1 and AH2, with offset hydrophobic faces. Both AH1 and AH2 are important for oligomerization of 2C in solution, with AH2 conferring most of the oligomerization propensity.

### The AH2 region mediates charge-independent binding of 2C to membranes

We next wanted to study the individual roles of AH1 and AH2 in 2C’s membrane binding. To this end, we reconstituted the process *in vitro* using synthetic liposomes, and the purified 2C constructs described above. We devised an assay in which MBP-fused 2C is first mixed with small or large unilamellar vesicles (SUVs and LUVs, respectively), after which TEV protease is added to cleave off the MBP tag and allow 2C to bind the liposomes (Fig. 3A, Fig. S5). The 2C-decorated liposomes are then subjected to a flotation assay, in which the liposome-containing upper fraction and liposome-free lower fraction are separated. 2C and liposomes are labeled with different fluorophores (ATTO488 maleimide conjugated to cysteines, and ATTO647N conjugated to DOPE, respectively), allowing quantification of 2C and lipids in the fractions (Fig. 3A, Fig. S6). 2C fluorophore labeling was performed on the native cysteines in its Cys-rich region containing a four-cysteine Zn finger. The two more solvent-exposed, and thus more likely to be labelled, cysteines in the Zn finger (C272 and C286) are known to be dispensable for poliovirus replication[43, 59]. We thus reasoned that one cysteine per monomer could probably be labeled without loss of Zn^2+^ coordination. This was confirmed by measurement of Zn^2+^ in labeled and unlabeled MBP-2C preps which showed no significant difference in Zn^2+^ content (Fig. S7). The lipid fluorescence readings in the upper fraction were consistent across all experiments, allowing for quantitative comparisons of the amount of protein bound to liposomes (Fig. S6).

**Figure 3:**
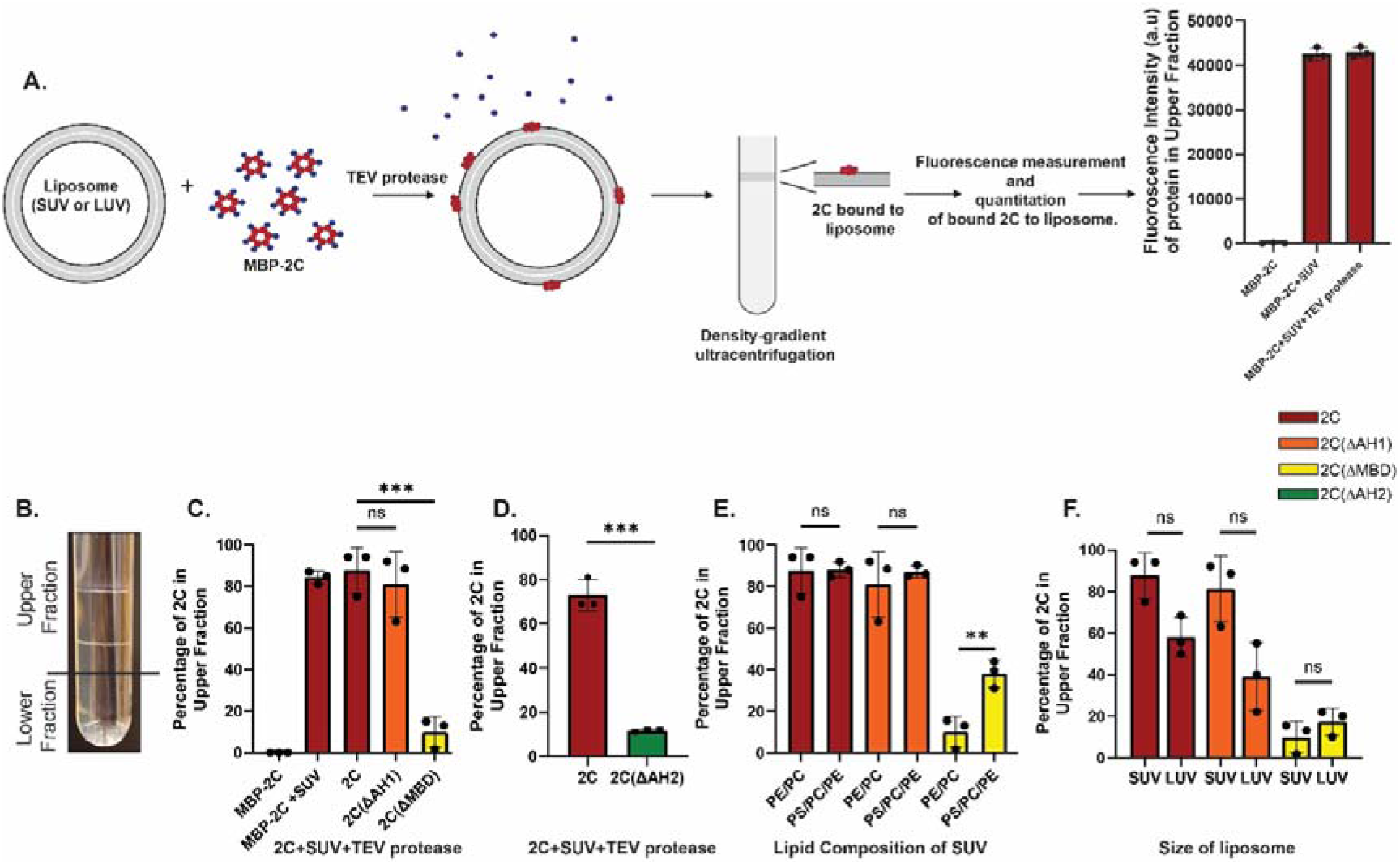
The AH2 region is critical for 2C’s membrane binding. **(A)** Schematic representation of flotation assay used to quantify 2C membrane binding. MBP-2C is fluorophore-labeled with ATTO488 on exposed cysteines, which only exist in the 2C part of the fusion protein. Small unilamellar vesicles (SUVs) or large unilamellar vesicles (LUVs) – collectively: liposomes – are labelled with DOPE-coupled ATTO647N. The MBP tag is cleaved from MBP-2C using TEV protease in the presence of liposomes, allowing the liberated MBD to directly associate with the liposomes. 2C-decorated liposomes are layered at the bottom of a density gradient and subjected to ultra-centrifugation. The amount of liposomes present in the top fraction, and their bound 2C, is quantified in separate fluorescence channels. 2C hexamers are dark red, MBP blue. **(B)** Photograph of a representative centrifuge tube after the flotation assay with 2C, showing two distinct lipid bands in the upper fraction. **(C-D)** Flotation assays conducted as in (A), with SUVs composed of 75 mol% POPC and 25 mol% POPE. 2C fluorescence was measured separately in upper and lower fractions, and the percentage present in the upper fraction is plotted for the different 2C constructs. **(E)** Flotation assay as in C, but with varying SUV lipid composition. PE/PC: 75 mol% POPC, 25 mol% POPE. PE/PC/PS: 25 mol% POPC, 25 mol% POPE, 50 mol% POPS. **(F)** Flotation assay as in C, but with varying liposome size. SUV: ∼50 nm diameter. LUV: ∼2500 nm diameter. **(C-F)** Error bars represent a standard deviation of three repeats of the experiment. Statistical significance by unpaired two-tailed Student’s t-test; ns: p>0.05, **: p<0.01, ***: p<0.001. Fig. S6 shows these data normalized by upper fraction lipid fluorescence.

We first used SUVs composed of net-uncharged lipids: 75 mol% 1-palmitoyl-2-oleoyl-glycero-3-phosphocholine (POPC) and 25 mol% 1-palmitoyl-2-oleoyl-sn-glycero-3-phosphoethanolamine (POPE). The SUVs had an average size of ∼30 nm as determined by dynamic light scattering (DLS) (Fig. S8). The addition of MBP-2C and TEV to the SUVs led to the formation of two distinct SUV bands in the upper fraction (Fig. 3B, Fig. S9A). 88±11% of full-length 2C were found in the upper fraction, with most protein in the uppermost of the two bands (Fig. S9B). MBP-fused 2C was able to associate with SUVs to a similar degree as cleaved 2C, indicating that the MBD is exposed in the MBP fusion protein. However, later experiments would show that removal of the MBP tag was necessary for functional association of 2C with membranes. Upon deletion of AH1, 81±16 % of 2C(ΔAH1) was still found in the upper fraction, whereas deletion of the entire MBD (AH1-AH2) led to only 10±7.4 % of 2C(ΔMBD) being found in the upper fraction (Fig. 3C). Further, the 2C(ΔAH2) construct at (11±0.6 %) had a similarly low membrane association as 2C(ΔMBD) (Fig. 3D). Thus, the AH2 region is the main mediator of 2C membrane association.

We then wished to study if 2C has a preference for a given membrane charge or curvature. In side-by-side flotation experiments, we quantitated 2C binding to the POPC/POPE (net uncharged) SUVs described above, and SUVs consisting of 50 mol% phosphatidyl serine (POPS, net charge -1). Full-length 2C and 2C(ΔAH1) showed no preference for the negatively charged POPS, whereas the binding of 2C(ΔMBD) increased from 10±7.4 % to 38±7% when POPS was included in the SUVs (Fig. 3E). To determine if 2C has a membrane curvature preference, we compared 2C binding to the sonicated POPC/POPE SUVs, with an average size of ∼30 nm, and larger (and hence lower-curvature) LUVs with the same lipid composition and an average diameter of ∼2500 nm. Full-length 2C and 2C(ΔAH1) showed a slight decrease in the binding to the lower-curvature LUVs (Fig. 3F), but this decrease was no longer significant after normalizing for the amount of lipid present in the top fraction (Fig. S6H).

In summary, these data show that the AH2 region mediates strong 2C membrane binding, which is neither dependent on high membrane curvature nor on negatively charged lipids.

### 2C mediates lipid clustering, which is enhanced by the AH1 region

In virus-infected cells, 2C is present at contact sites of ROs and lipid droplets[24], and we suspected that the high-density band of 2C-coated SUVs in our flotation experiments may be an *in vitro* recapitulation of 2C-mediated lipid clustering (Fig. 3B, Fig. S9A). We used DLS to investigate lipid clustering by 2C. As described above, the MBP tag was cleaved from 2C in the presence of SUVs. After a 30 min incubation, the size distribution of the sample was measured using DLS. The sonicated SUVs were around 40 nm in size, whereas the addition of TEV protease and 0.8 µM MBP-2C to these SUVs led to the formation of vesicle clusters with an average diameter of ∼1 μm (Fig. 4A). 2C(ΔAH1) also clustered SUVs (Fig. 4A). Defining a liposome cluster as a particle with a diameter of >100 nm (2.5 times the average SUV size), we compared the percentage of clusters in reactions containing different concentrations of 2C and 2C(ΔAH1). The lack of binding to PC/PE membranes by 2C(ΔMBD) in the flotation assay meant that we could rule out that it would cluster membranes (an activity which requires binding of two membranes) and it was thus not studied in this assay. Already at 0.8 μM protein – the concentration used for the flotation assays – full-length 2C generated significantly more clusters than 2C(ΔAH1) (11.5±4.0% and 4.6±1.7%, respectively, Fig. 4B). At 3 μM 2C, the difference was even more pronounced, with 47.0±13% of liposomes being present in clusters with 2C and 12±13% for 2C(ΔAH1) (Fig. 4B). To validate our findings, we performed negative-staining TEM on identical reactions to those studied by DLS. The trend was similar to DLS, i.e. relatively more clustering with full-length 2C as compared to 2C(ΔAH1), and no visible clustering for 2C(ΔMBD) and 2C(ΔAH2) (Fig. S10). We then performed cryo-electron tomography (cryo-ET) on the 3 μM reaction with full-length 2C. This allowed direct visualization of 2C tethering liposomes together to form larger clusters (Fig. 4C). Typically, multiple protein densities were present on the imaged SUVs, mostly at the interface between SUVs (Fig. 4C). Based on the volumes of individual densities, their masses were estimated to 199±53 kDa, closely matching the theoretical mass of a 2C hexamer at 222 kDa (Fig. 4D). Taken together, 2C clusters liposomes *in vitro*, and the AH1 region is necessary for effective clustering.

**Figure 4:**
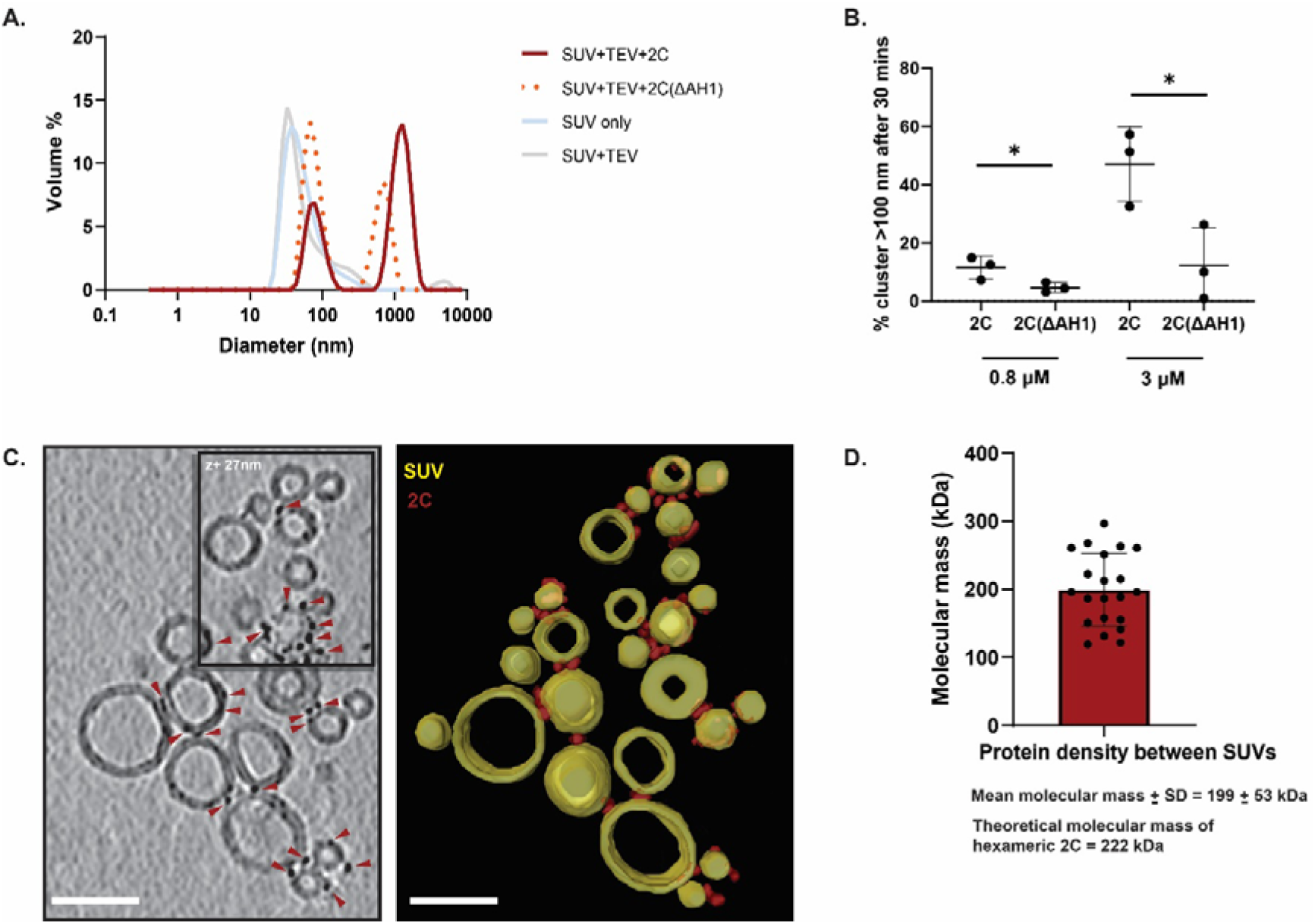
2C has lipid clustering activity that is enhanced by the AH1 region. **(A)** Representative dynamic light scattering (DLS) graphs (size distribution by volume), showing the size distribution of the PE/PC SUV preparation, and appearance of additional peaks corresponding to vesicle clusters, 30 min after adding TEV protease and 3 μΜ MBP-2C (red solid line) or MBP-2C(ΔAH1) (orange dotted line) to SUVs (blue solid line). **(B)** Percentage of the total vesicle volume present in clusters (defined as particles with diameter > 100 nm), 30 min after adding TEV protease and 0.8 or 3 µM MBP-2C or MBP-2C(ΔAH1). Dots represent the average of triplicate measurements on the same SUV preparation, bars the average and error bars one standard deviation of the three independent replicates. Statistical significance by unpaired two-tailed Student’s t-test; ns: p>0.05, **: p<0.01 **(C)** Computational slice through a representative cryo-electron tomogram of PE/PC SUVs, 30 min after adding TEV protease and 3 μΜ MBP-2C. Densities corresponding to 2C are indicated by red triangles. The right panel contains the corresponding three-dimensional segmentation showing 2C (red) and SUVs (yellow). Scale bar, 50 nm. **(D)** Molecular mass estimation based on volume of individual densities found between the SUVs. Each dot represent one density. The molecular mass was estimated to be 199±53 kDa (mean±std).

### Nucleic acid binding modulates 2C membrane clustering

The enterovirus genome is replicated on the cytoplasmic face of the RO membrane [20, 60]. Since 2C is a AAA+ protein and a putative RNA helicase [35, 38, 46, 48, 49, 61–64], we wondered whether 2C is sufficient to recruit and retain RNA at the membrane, and whether nucleic acid binding may affect 2C’s lipid clustering activity. To address the second question, we repeated the lipid clustering experiment using full-length 2C, in the presence of increasing concentrations of nucleic acid. It was previously reported that both DNA and RNA bind at the same site on 2C [48]. Hence, we used single-stranded DNA (ssDNA) oligonucleotides for the initial experiment, and measured the amount of clustering using DLS (Fig. 5A). Upon increasing the concentration of ssDNA there was a consistent decrease in the percentage of clustered lipids, albeit only statistically significant at 100 μM ssDNA concentration (Fig. 5B). In a comparison performed at a lower concentration (5 µM), ssRNA had a stronger effect on modulation of clustering than ssDNA (Fig. 5C). This indicates that nucleic acid binding partially outcompetes the membrane tethering activity of 2C. To directly test the hypothesis that 2C recruits RNA to the membrane, and to investigate if it possesses any selectivity, we devised an alternative SUV flotation assay. In this assay, 2C was no longer fluorophore-labelled. Instead, a fluorophore-labeled RNA (single-stranded or double-stranded) was added in the first step of the flotation assay, allowing quantitation of the amount of RNA co-migrating with the SUVs (Fig. 5D, Fig. S11). 2C was able to recruit both ssRNA and dsRNA to SUVs and retain some of it at the membrane throughout the flotation experiment (Fig. 5E). At identical starting concentrations, dsRNA was recruited significantly more than ssRNA (Fig. 5E). The recruitment was strictly dependent on 2C having a free MBD since omission of TEV protease completely abrogated the RNA recruitment (Fig. 5E). Taken together, 2C recruits and retains nucleic acid to membranes, with a preference for dsRNA over ssRNA, and nucleic acid binding reduces 2C-mediated membrane clustering.

**Figure 5:**
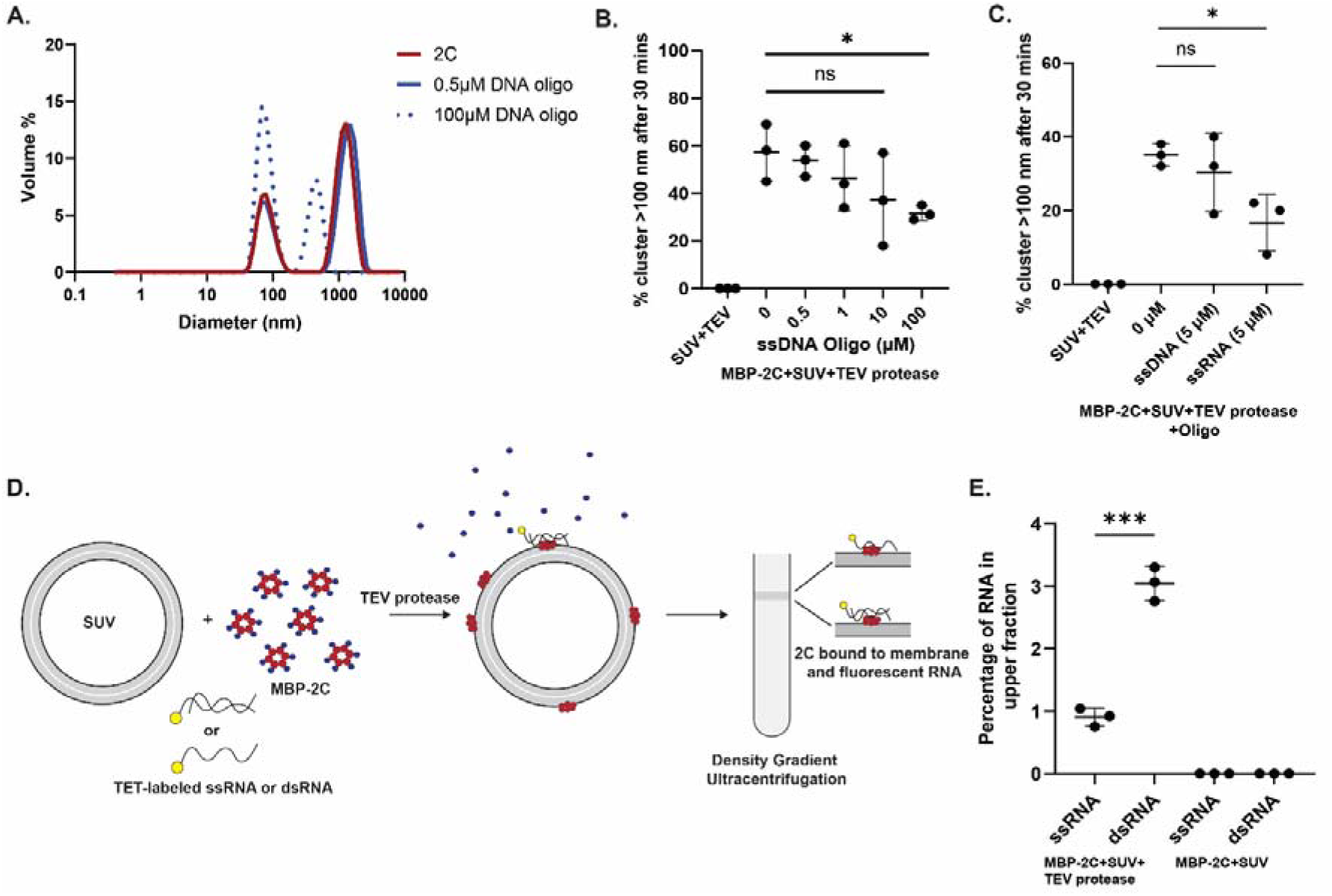
Nucleic acid binding modulates 2C membrane clustering. **(A)** Representative DLS graphs (size distribution by volume), performed as for Fig. 4A with 3 μM 2C, but with the indicated concentrations of ssDNA oligonucleotide added. **(B)** Percentage of vesicle volume present as clusters (evaluated as for Fig. 4B), 30 min after addition of TEV protease, MBP-2C, and the indicated concentration of ssDNA oligonucleotide. **(C)** Same assay as in (B) except that 5 µM ssDNA or ssRNA was added. **(D)** Schematic representation of the flotation assay used to quantify RNA recruitment to membranes by 2C. The experiment is essentially performed as in Fig. 3A, except that MBP-2C is unlabeled, and the RNA (single-stranded, or annealed with a complementary strand for double-stranded) is labeled with TET fluorophore at its 5′ end (Fig. S11). Membrane-bound RNA is quantitated as TET fluorescence in the upper fraction **(E)** Fluorescence from ssRNA and dsRNA in top fraction in the experiment outlined in (C). **(B,C,E)** Each dot represent the average of three measurements on one SUV preparation, bars the average and error bars one standard deviation of the three independent replicates. Statistical significance by unpaired two-tailed Student’s t-test; ns: p>0.05, *: p<0.05, ***: p< 0.001.

### Membrane-bound, full-length 2C is an ATPase with single-strand ribonuclease activity

Several studies have sought to determine whether 2C is an active RNA helicase, but they were generally conducted with 2C that was truncated and/or not membrane-associated. Having reconstituted the binding of full-length 2C to membranes, and shown that it recruits RNA to membranes, we set out to use this more physiologically relevant system to revisit 2C’s helicase activity. The membrane association of 2C did not alter its ATPase activity as compared to MBP-2C in solution (Fig. 6A). We then conducted an RNA helicase assay with membrane-bound 2C. The unwinding of dsRNA to ssRNA was monitored on a native polyacrylamide gel. Due to preliminary observations that TEV protease can affect the helicase assay, its concentration was decreased until it no longer affected the migration of the double-stranded RNA substrate (Fig. 6B). Under these conditions, a substantial, albeit not stoichiometric, fraction of MBP was released from MBP-2C during the assay (Fig. S12). With dsRNA substrate having a single-strand 5′ overhang, a clear ATP-dependent unwinding was observed for the positive control Pfh1, a well-characterized, 5′ overhang-specific *S. pombe* helicase (Fig. 6B-C) [65, 66]. However, in the case of membrane-bound 2C, the addition of ATP and Mg^2+^ did not result in formation of single-strand RNA product of the expected size (Fig. 6B-C). Instead, lower molecular-weight bands, corresponding to cleaved fluorophore-labelled RNA, appeared (Fig. 6B,D). The appearance of the cleaved products was dependent on Mg^2+^ but independent of ATP (Fig. 6E, Fig. S13A). Notably, all assays were conducted using reaction components prepared in DEPC-treated water and in the presence of RNase inhibitors. We then repeated the same experiment with dsRNA having a 3′ overhang (Fig. 6F). Due to its specificity for 5′ overhangs, the control helicase Pfh1 was not included in this assay. Analogously to the 5′ overhang substrate, no helicase activity was detected with the 3′ overhang substrate (Fig. 6F-G). On the other hand, the ribonuclease activity was even more pronounced with the 3′ overhang substrate (Fig. 6F,H), and again shown to be dependent on Mg^2+^ but independent of ATP (Fig. 6I, Fig. S13B). The ribonuclease activity of 2C was comparable when using fluorophore-labeled single-strand RNA as substrate (Fig. S13C-D). In summary, full-length, membrane-bound 2C is an active ATPase and has Mg^2+^-dependent ribonuclease activity, but no RNA helicase activity under the conditions of this study.

**Figure 6:**
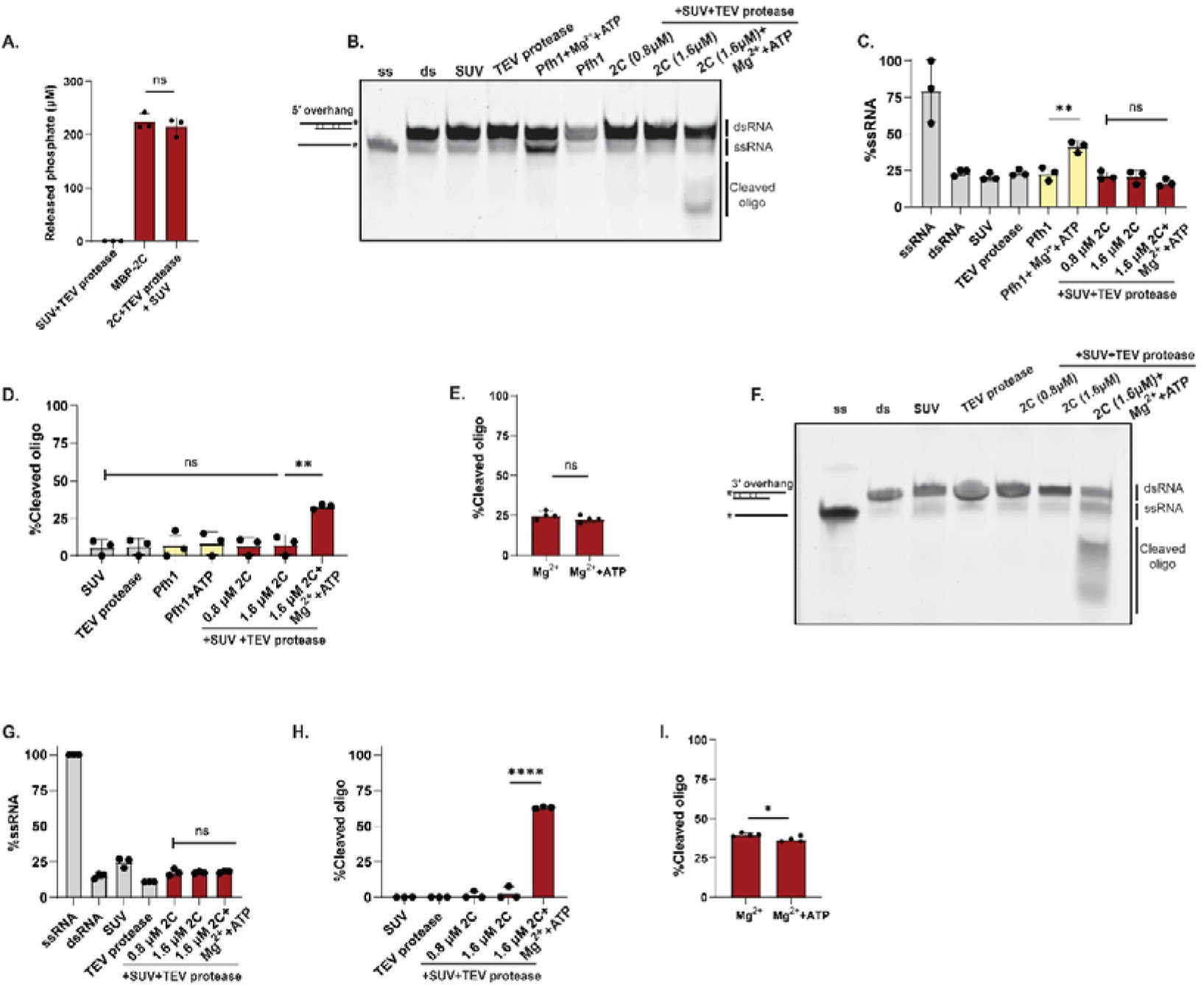
Membrane-bound 2C has ATPase and ribonuclease activity. **(A)** Colorimetric ATPase assay with MBP-2C, 0.4 mM ATP and other components as indicated. **(B)** Fluorescence scan of native polyacrylamide gel resolving dsRNA and ssRNA. Lanes 1 and 2 from the left were loaded with ssRNA control and dsRNA substrate (5′ overhang and fluorophore on 3′ end), respectively. The remaining lanes were loaded with reactions taken at the end of the incubation with dsRNA substrate other components as indicated. Pfh1, positive control helicase. A representative gel is shown, the experiment was performed in three independent replicates. **(C)** The percentage of total RNA fluorescence present as ssRNA for selected lanes of the gel in (A), and two independent replicates of the same experiment. **(D)** Gel quantitation as in (B) but for cleaved oligo. **(E)** Quantitation of the gel shown in Fig. S13A and two independent replicates. The assay conditions are the same as in the rightmost bar in (C), except when Mg^2+^ was added without ATP. **(F)** Fluorescence scan of a gel for an experiment similar to (A), but with a 3′ overhang on the dsRNA (fluorophore on 5′ end). **(G-H)** Quantifications of the gel in (F) and two independent replicates, as in (C-D). **(I)** Quantitation of the gel shown in Fig. S13B, as in (E). **(A-I)** Dots represent individual experiments, bars the average and error bars one standard deviation. Statistical significance by unpaired two-tailed Student’s t-test; ns: p>0.05, *: p<0.05, **: p<0.01, ****: p<0.0001

### AH1 contributes to viral RNA replication to a similar degree as the entire MBD

The biochemical reconstitution allowed us to study the separate contributions of AH1 and AH2 to processes such as membrane binding and membrane clustering. To investigate their relative contribution to viral RNA replication in cells, we used poliovirus replicons where the capsid proteins were replaced by Renilla luciferase. As opposed to the biochemical experiments, the 2C truncations in replicons (Δ(2-11) and Δ(2-40)) retained the starting glycine of 2C since it is a core part of the viral protease 3C cleavage site (Fig. 7A). 12 hours after transfecting replicon RNAs into cells, they were harvested, and the luciferase signal was quantified. The luminescence from both the Δ(2-11) and Δ(2-40) replicons was reduced by approximately two orders of magnitude with respect to wildtype, a similar reduction to a control replicon with a catalytically inactive 3D polymerase (Fig. 7B). Thus, although AH1 appeared to have more subtle roles in the biochemical experiments (e.g. in vesicle clustering), it is indispensable for effective viral RNA replication in cells.

**Figure 7:**
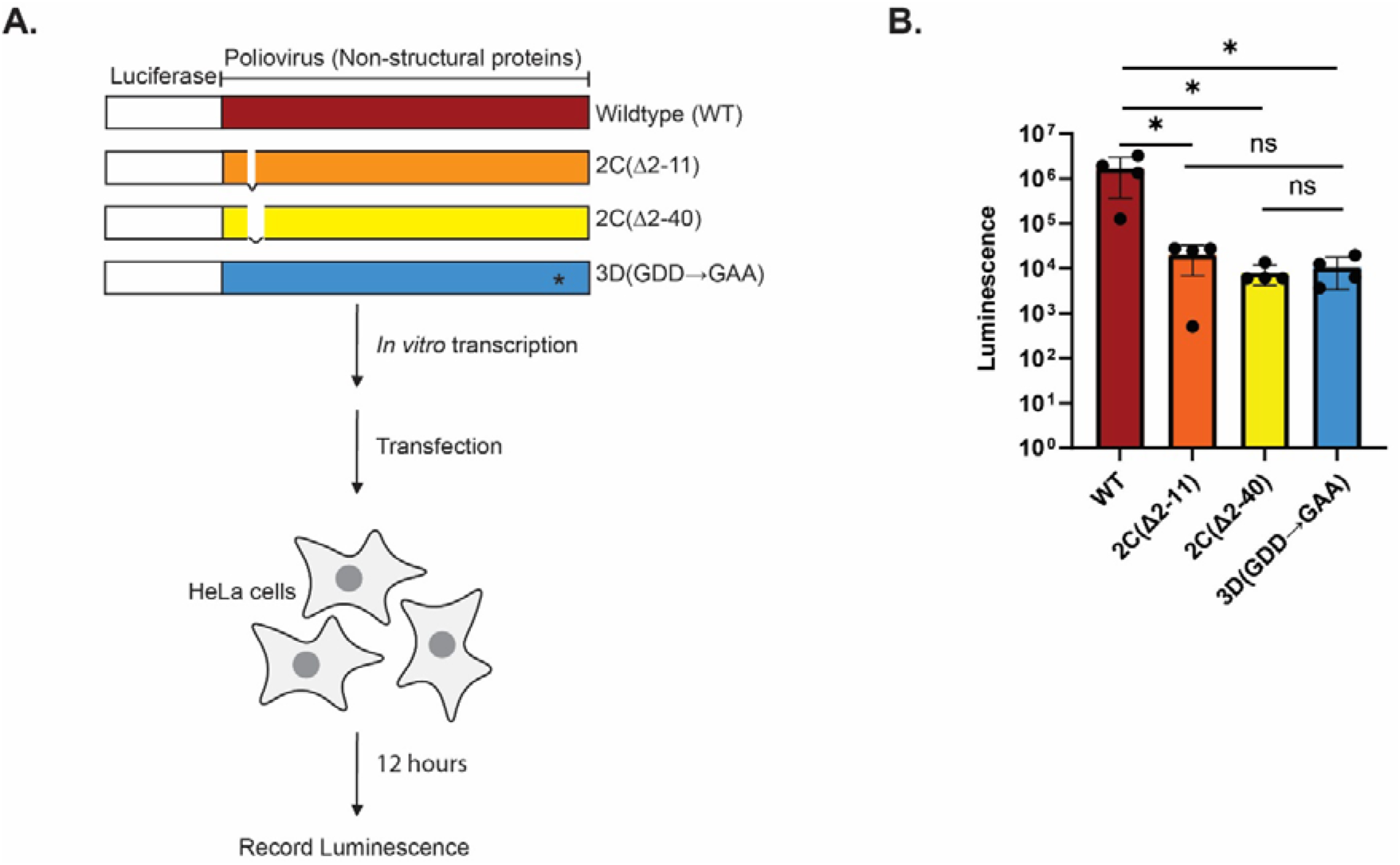
Truncation of AH1 or the MBD drastically reduces poliovirus genome replication. **(A)** Schematic representation of poliovirus replicons and their use. The replicons contained Renilla luciferase in place of the capsid region of the polyprotein, and the indicated deletions in the MBD of 2C, or replication-defective 3D polymerase with the active site GDD-to-GAA change. The first residue of 2C, glycine, was kept in both deletions since it is a core part of the protease cleavage site. In vitro-transcribed RNAs from these replicons were transfected into HeLa cells and luminescence was recorded 12 hours post-transfection. **(B)** Luminescence recorded for cells transfected with different poliovirus replicons. Dots represent individual experiments, bars the average, and error bars one standard deviation of the four independent replicates. Statistical significance by unpaired two-tailed Student’s t test; ns p>0.05; *p < 0.05

## Discussion

With fewer than 7,000 coding nucleotides, the *Enterovirus* genome can sustain a full viral replication cycle by encoding multifunctional proteins, and by hijacking host-cell functions. A universally present and highly conserved enteroviral protein is the AAA+ ATPase 2C, whose suggested roles range from membrane remodeling to RNA helicase and capsid morphogenesis. The nature of 2C as an oligomerizing, peripheral membrane protein has impeded biochemical studies, which have typically been conducted on truncated protein not engaging a membrane. Here we described the *in vitro* reconstitution of purified, full-length 2C’s binding to membranes. We used this more physiologically relevant, yet completely defined, system to investigate several functions of 2C. In summary, our findings are consistent with the model shown in Fig. 8.

**Figure 8:**
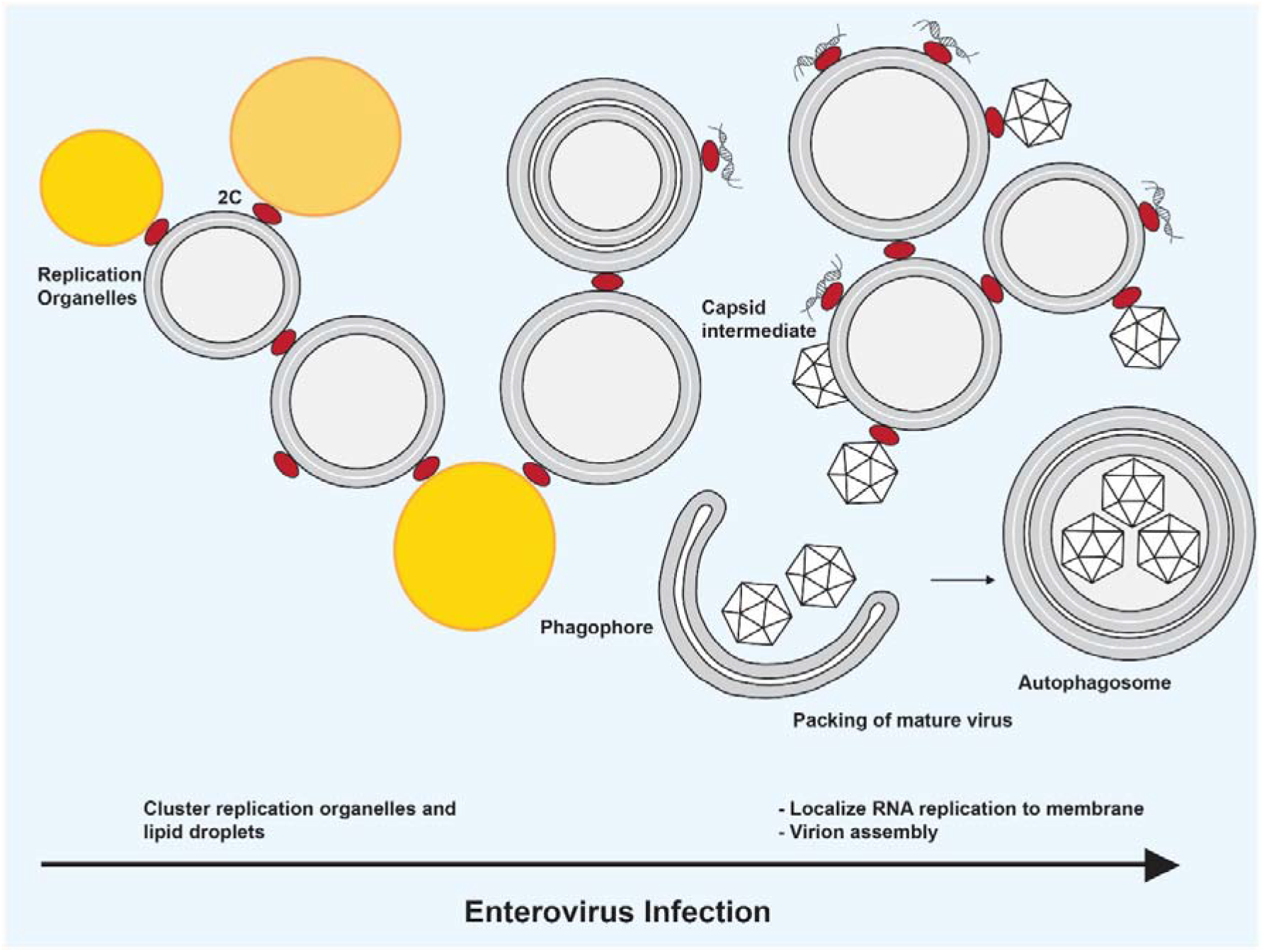
A model for the some of the roles of 2C in enterovirus replication. The model places the biochemical findings of this study into a suggested cellular context, which may include more roles of 2C and distinct roles for the partially cleaved polyprotein 2BC (not shown in the figure). At an early stage of infection, 2C (and 2BC) serves to cluster replication organelle membranes and lipid droplets, while tethering the first viral dsRNA to the membrane. As the replication progresses, 2C would be increasingly loaded with viral dsRNA and thus play a role in localizing viral RNA replication to the replication organelle membranes. How, or if, this mechanism leads to the shielding of the viral dsRNA from innate immune sensors is not yet known. An increasing amount of 2C in the course of the infection means that a fraction of 2C may still be involved in e.g. membrane clustering in late infection. Our data suggest that 2C is an active ATPase but not an RNA helicase – a role which may instead be played by the polymerase 3D[74]. Although not addressed in this study, the literature strongly suggests an additional role for 2C in capsid assembly and RNA loading, consistent with it being part of a “tethering complex” which tethers newly synthesized capsids to replication organelle membranes[20].

We integrated various structure prediction methods to conclude that the N-terminal region of 2C, which had previously been identified as its amphipathic, membrane-binding domain (MBD)[50–54], is in fact divided into two amphipathic helix segments. The two amphipathic segments, AH1 and AH2, are separated by a highly conserved glycine residue (Fig. 1B-C, Fig. S1). In the longer term, experimental structures of full-length, membrane-bound 2C are needed to test these and other predictions about 2C’s structure. Such structures may be achievable by cryo-electron tomography and subtomogram averaging[67, 68]. The second predicted amphipathic helix segment, AH2, is the main mediator of 2C’s oligomerization in solution (Fig. 2), and is sufficient for membrane binding (Fig. 3C). This finding aligns well with the observation that an isolated peptide corresponding to AH2 is able to bind liposomes[69]. Partial membrane binding of 2C(ΔMBD) could be restored by including 50% POPS in the liposomes, which indicates that 2C has a second membrane-interacting site, which is dependent on membrane charge, outside of the MDB (Fig. 3E). Of note, removal of the N-terminal maltose-binding protein (MBP) tag was not necessary for 2C to bind membranes in our assay, but it was strictly required for 2C to recruit RNA to membranes (Fig. 5E). Whereas the N-terminal MBP tag is an artificial addition to aid purification of recombinant 2C, the fusion of 2C’s N-terminus to another protein is, in fact, relevant to infection. 2C is initially excised from the viral polyprotein in the form of the metastable precursor 2BC, in which the N terminus of 2C is still fused to the viral protein 2B [70]. Hence, our finding that 2C only binds RNA after its N terminus is liberated may be indicative of what happens in an infected cell when 2BC is processed to 2B and 2C. Ultimately, this hypothesis would need to be tested in an experimental system including native 2BC and its cleavage by the viral protease 3C, since the properties of 2B are significantly different from those of MBP.

Our study revealed a novel role for 2C in clustering vesicles, a role mediated by AH1 (Fig. 4). This finding suggests a biochemical mechanism for the 2C-mediated recruitment of lipid droplets to replication organelles (ROs)[24]. 2C-mediated membrane clustering may also be involved in generating the tightly stacked membrane tubes that make up the ROs at early time points [21, 71]. Notably, the deletion of AH1 led to a stark reduction of poliovirus genome replication, similar in effect to the deletion of the entire MBD (Fig. 7). This may indicate that the lipid-clustering activity of 2C is central to enterovirus replication, although it cannot be ruled out that AH1 deletion also affects other aspects of the replication such as polyprotein processing[72]. The decrease in 2C-mediated membrane clustering seen with increasing nucleic acid concentration (Fig. 5A-C) suggests a balance between 2C’s role in membrane clustering at early time points, and a preferential role in RNA replication and particle assembly at later time points (Fig. 8). In this context, the ability of 2C to bind membranes independent of curvature and lipid species (Fig. 3E-F) fits its multitude of roles in binding to different types of RO membranes and lipid droplets.

The localization of viral RNA replication to RO membranes is well established, but the molecular mechanism behind this localization is poorly understood. An electrostatic interaction between the viral polymerase 3D and anionic lipids may be part of the mechanism[73]. Here, we show that 2C is able to recruit and retain RNA at membranes, with a preference for dsRNA (Fig. 5E). Long double-stranded segments are not prevalent amongst cellular RNAs, but it is the form of the viral template RNA while replicated. Thus, 2C could selectively localize viral template RNA to the ROs. On the other hand, full-length, membrane-bound 2C had no detectable RNA helicase activity under the conditions of our assay (Fig. 6). This is in line with many previous studies on truncated and membrane-free 2C, and with structural models of the 2C hexamer indicating that its central pore would be too narrow and have an electrostatic potential incompatible with RNA translocation [44, 45, 48]. The convergence of our work and previous studies in suggesting that 2C is not a helicase can be interpreted in the light of the reported dsRNA-unwinding activity of the polymerase 3D [74]. However, it remains a possibility that 2C has RNA helicase activity specific to certain RNA secondary structures, which has been demonstrated for some other helicases[65, 75]. It is also a possibility that post-translational modifications, which are not recapitulated upon bacterial protein expression, activate 2C has a helicase[49]. In our assay designed as a helicase assay, we discovered that membrane-bound 2C has ribonuclease activity on dsRNA with single-strand overhangs, as well as on ssRNA (Fig. 6). The activity was present on substrates having both a 3′ and 5′ fluorophore label, thus ruling out a direction-specific exonuclease activity. A recent paper reported single-stranded endonuclease activity of truncated and non-membrane-bound 2C from hepatitis A virus as well as several enteroviruses[76]. Our work shows that this activity also exists in the context of full-length 2C on a membrane, which warrants further investigation into its role in virus replication.

Taken together, our reconstitution of 2C membrane binding allowed us to shed light on several functions of 2C: it is able to bind vesicles with no apparent preference for curvature and lipid composition, it can cluster vesicles, selectively recruit dsRNA to membranes and function as a ribonuclease. In spite of this, major questions remain unanswered about the role of 2C in enterovirus RNA replication and particle assembly. Nothing is known about the supramolecular arrangement of the RNA replication machinery on membranes, for instance how 2C, the polymerase 3D and host RNA-binding proteins such as PCBP1/2 are arranged [77]. Even though we show that 2C can bind dsRNA, the mechanism of shielding of dsRNA from innate immune sensors is not known. Also, 2C is known to play a role in capsid assembly and is suggested to be part of a “tether complex” enabling the RNA loading of new capsids on RO membranes [20, 71], but biochemical mechanisms underlying this process are largely unknown. The reconstitution of 2C-decorated vesicles presented here is a stepping stone towards a more complete *in vitro* reconstitution of such complex processes.

## Materials and methods

### Expression and purification of poliovirus 2C

A codon-optimized synthetic gene corresponding to full-length 2C from poliovirus type 1 Mahoney (GenBank: V01148.1) was inserted into 1M expression plasmid from Macrolab (University of California, Berkeley, USA) with a TEV protease-cleavable, N-terminal His_6_-MBP tag. The plasmid was transformed to *E. coli* BL21(DE3) competent cells and grown overnight on LB-agar plates containing 50μg/ml Kanamycin (KAN). A single colony was used to inoculate LB/KAN media, this primary culture was grown for 5 hours. Then, a one-liter culture of LB/KAN was inoculated with 1ml of the primary culture. The cells were grown at 37°C, induced at OD_600_=2 by the addition of 0.5 mM isopropyl-beta-D-thiogalacopyranoside, then incubated overnight at 25°C. Cells were collected by centrifugation, and resuspended in lysis buffer (50 mM Tris, pH 8.5), 500 mM NaCl, 20 mM Imidazole, 10% (v/v) glycerol, 36 μM NP40, 5 mM MgCl_2_, 0.1 mM THP) containing protease inhibitor cocktail (40 mM Phenylmethylsulfonyl fluoride, 1 mM Leupeptin and 200 mM Benzamidine HCl) and DNase. The cells were lysed using a continuous flow cell disruptor (Constant Systems). The lysate was cleared by centrifugation at 63,000*g at 4°C for 1 hour and the supernatant was loaded onto Ni-NTA (nitrilotriacetic acid) resin and eluted with elution buffer (50 mM Tris (pH 8.5), 500 mM NaCl, 250 mM Imidazole, 10% (v/v) glycerol, 36 μM NP40, 5mM MgCl_2_, 0.1 mM THP). The protein was then loaded onto a HiTrap Q HP Column (GE Healthcare) and eluted using a gradient from buffer QA (50 mM Tris, pH 8.5), 50 mM NaCl, 10% (v/v) glycerol, 36 μM NP40, 5 mM MgCl_2_, 0.1 mM THP) and buffer QB (50 mM Tris (pH 8.5), 1 M NaCl, 10% (v/v) glycerol, 36 μM NP40, 5 mM MgCl_2_, 0.1 mM THP). The fractions containing the protein was loaded onto a 24 ml Superose 6 Increase 10/300 GL (Cytiva) that was equilibrated with SEC buffer (50 mM Tris-HCl (pH 7.4), 150 mM potassium acetate, and 0.1 mM THP). Purified 2C was then aliquoted, flash frozen with liquid N_2_, and stored at -80°C. All the other constructs of the protein were purified in a similar way.

To prepare 2C(41-329) and 2C(116-329) without His_6_-MBP tag, the tagged proteins were diluted to 0.1mg/ml after the HiTrap Q HP column using buffer QA described above and the proteins were cleaved at 4°C overnight by adding TEV protease in a molar ration TEV:2C of 1: 5 and 1:15 for 2C(41-329) and 2C(116-329), respectively. Uncleaved protein and His_6_-MBP were removed by incubating the mixture with NiNTA resin followed by passing of the protein over a 1ml MBPTrap (Cytiva) column and then a HiTrap Heparin HP (Cytiva) using buffer QA and QB. The fractions containing the protein were loaded onto a Superdex 75 Increase 10/300 GL (Cytiva) equilibrated with SEC buffer. Purified cleaved 2C was aliquoted, flash frozen with liquid N_2_, and stored at -80°C.

### Fluorophore labeling of proteins

To label the proteins, after the HiTrap Q HP Column, the proteins were incubated with an equimolar amount of Atto488 maleimide (Sigma-Aldrich) at 4°C overnight. Next day, excess dye was quenched using 25 mM Dithiothreitol (DTT) and the proteins were loaded onto a Superose 6 Increase 10/300 GL (Cytiva) column, pooled and snap frozen as described above for unlabeled protein. The labeling efficiency was estimated by measuring the absorption of the labeled protein at 280 nm and at the fluorophore absorption maximum. It was typically 0.5-1.1 fluorophores/2C monomer.

## SEC-MALS

All the samples were run on an ÄKTA Pure (Cytiva) with a Superdex 200 Increase 10/300 GL column, equilibrated in SEC buffer, coupled to a light scattering (Wyatt Treas II) and refractive index (Wyatt Optilab T-Rex) detector. Bovine serum albumin (BSA) was used to calibrate the system. 150 to 500 µl (0.6 to 1 mg/ml) of protein was injected at a flow rate of 400 µl/min. Data were analyzed using Astra software (version 7.2.2; Wyatt Technology).

### Mass photometry

Proteins were diluted to 100 nM in 50 mM Tris pH 7.4 and applied to a glass coverslip and recorded on Mass Photometer[78] (OneMP, Refeyn Ltd) for 60 seconds. The data were analyzed using DiscoverMP (Refeyn Ltd) and the molecular weight was obtained by contrast comparison with known mass standards measured on the same day and were repeated three times, and visualized as mean +/-standard deviation.

### Multiple sequence alignment

Multiple sequence alignment of amino acids 1-40 was performed on EV71 (GenBank: AAB39969.1), CVB3 (GenBank: AFD33642.1), PV type 1 (MAHONEY STRAIN) (GenBank: V01148.1), EV D68 (GenBank: ALE66301.1), RV A2 STRAIN USA/2018/CA-RGDS-1062 (GenBank: MN228695.1) and RV C15 (GenBank: ACZ67658.1) using Clustal Omega[79] with default parameters.

### Amphipathic helix prediction

The physicochemical properties 2C amino acids 1-40 from EV71 (GenBank: AAB39969.1), CVB3 (GenBank: AFD33642.1), PV type 1 (Mahoney strain) (GenBank: V01148.1), EV D68 (GenBank: ALE66301.1), RV A2 STRAIN USA/2018/CA-RGDS-1062 (GenBank: MN228695.1) and RV C15 (GenBank: ACZ67658.1) were determined using heliquest (https://heliquest.ipmc.cnrs.fr/) [80] using alpha-helix as helix type and window size of one turn (18 residues). The mean amphipathic moment or transverse hydrophobic moment (µH) was plotted against the amino acid at the center of the analysis window.

### Structure predictions

All structures were predicted using Alphafold 2 implemented in COLABFOLD ((https://colab.research.google.com/github/sokrypton/ColabFold/blob/main/AlphaFold2.ipynb) [55] with default settings. The cartoon representations were generated using CHIMERAX [56].

### Preparation of small unilamellar vesicles (SUVs)

1-palmitoyl-2-oleoyl-sn-glycero-3-phospho-L-serine (POPS), 1-palmitoyl-2-oleoyl-sn-glycero-3-phosphoethanolamine (POPE) and 1-palmitoyl-2-oleoyl-sn-glycero-3-phosphocholine (POPC) were purchased from Avanti Polar lipids. Total 1 mg/ml of lipids containing POPC (25 mol %), POPE (25 mol %), POPS (25 mol %) or POPC (75 mol %), POPE (25 mol %), and 1, 2-Dioleoyl-sn-glycero-3-phosphoethanolamine labeled with Atto 647N (0.1 mol %) was dried onto a borosilicate glass tube wall with a nitrogen stream and kept overnight in vacuum desiccator for complete removal of solvents. 500 µl of SEC buffer was added into the mixture. The tube was then vortexed to resuspend the lipids and then, kept in a water bath at 42°C for 15 min. Then, the lipids were transferred to an Eppendorf tube for sonicating with a tip sonicator (10 seconds pulse, 20 seconds rest, and 20% amplitude) until a clear solution was observed. Then, the tube was centrifuged at 21,000*g at 4°C for 20 min. Then, 300 µl of supernatant was carefully removed, and the quality of the preparation was checked using dynamic light scattering as described below. SUVs were kept at 4°C until use, at most for 3 days.

### Preparation of large unilamellar vesicles (LUVs)

Total 1 mg/ml of lipids containing POPC (75 mol %), POPE (24.9 mol %) and1, 2-Dioleoyl-sn-glycero-3-phosphoethanolamine labeled with Atto 647N (0.1 mol %) was dried onto a borosilicate glass tube wall with a nitrogen stream and kept overnight in vacuum desiccator. Then, 500 µl of SEC buffer was added into the mixture. The tube was then vortexed to resuspend the lipids and to break the multi-lamellar vesicles, the lipids were subjected to 12 freeze-thaw cycles by moving it between a 42°C water bath and liquid nitrogen. Then, this solution was passed 21 times through an extrusion membrane (Whatman Nuclepore) with pore size of 2 μm on a mini-extruder (Avanti Polar Lipids) to get a clear solution comprising of LUVs. The quality of the preparation was checked using Dynamic light scattering as described below. LUVs were kept at 4°C until use, at most for 3 days.

### Vesicle flotation assay

A reaction mixture of 100 µl was prepared containing 0.8 μM Atto488 maleimide His_6_MBP-2C, 1 mg/ml SUVs or LUVs and 8 μM TEV protease was incubated for 2 hours at room temperature. Then, this reaction mixture was mixed with 500 μl of 48% (w/v) OptiPrep density gradient medium (Merck (Sigma-Aldrich)) and this mixture was carefully placed at the bottom of an Open-Top Thin wall Ultra-Clear Tube (Beckman Coulter) and was overlaid with 800 μl of 30% (w/v) OptiPrep followed by another layer of 800 μl of 20% (w/v) OptiPrep and finally 2 ml of SEC buffer. The tubes were then ultra-centrifuged at 364,000*g for 3 hours at 4°C. After centrifugation different fractions were carefully taken out from the tubes and fluorescence was measured using Agilent BioTek Synergy fluorimeter.

### Flotation assay with fluorescently labeled RNA

All oligonucleotides used in this study are listed in Table 1. 100 μl of reaction mixture was prepared as described above, except that unlabeled 2C was used, and a final concentration of 500 nM of single-stranded 5′ TET-labeled RNA(5′CCUCUAACCACAGUCUGAUC 3′) (Eurofins Genomics) or dsRNA (described below) was added. The flotation assay was performed as described above and the TET fluorescence was measured. To prepare dsRNA, 10uM of 5′ TET labeled RNA (5′CCUCUAACCACAGUCUGAUC 3′) was mixed with its 10uM unlabeled complementary strand in a buffer containing 50mM HEPES pH7.2, 100mM KCl and RNase secure. The mixture was heated to 95°C and then the temperature was decreased by 0.5°C /minute until 22°C. The annealed dsRNA was then placed overnight at 4°C and then at -20°C until use. The extent of annealing was checked using a 16% acrylamide non-denaturing RNA gel. A standard curve was prepared using fluorescence reading for ssRNA and dsRNA and the extent of binding of RNA to 2C was quantified using this standard curve.

**Table 1:**
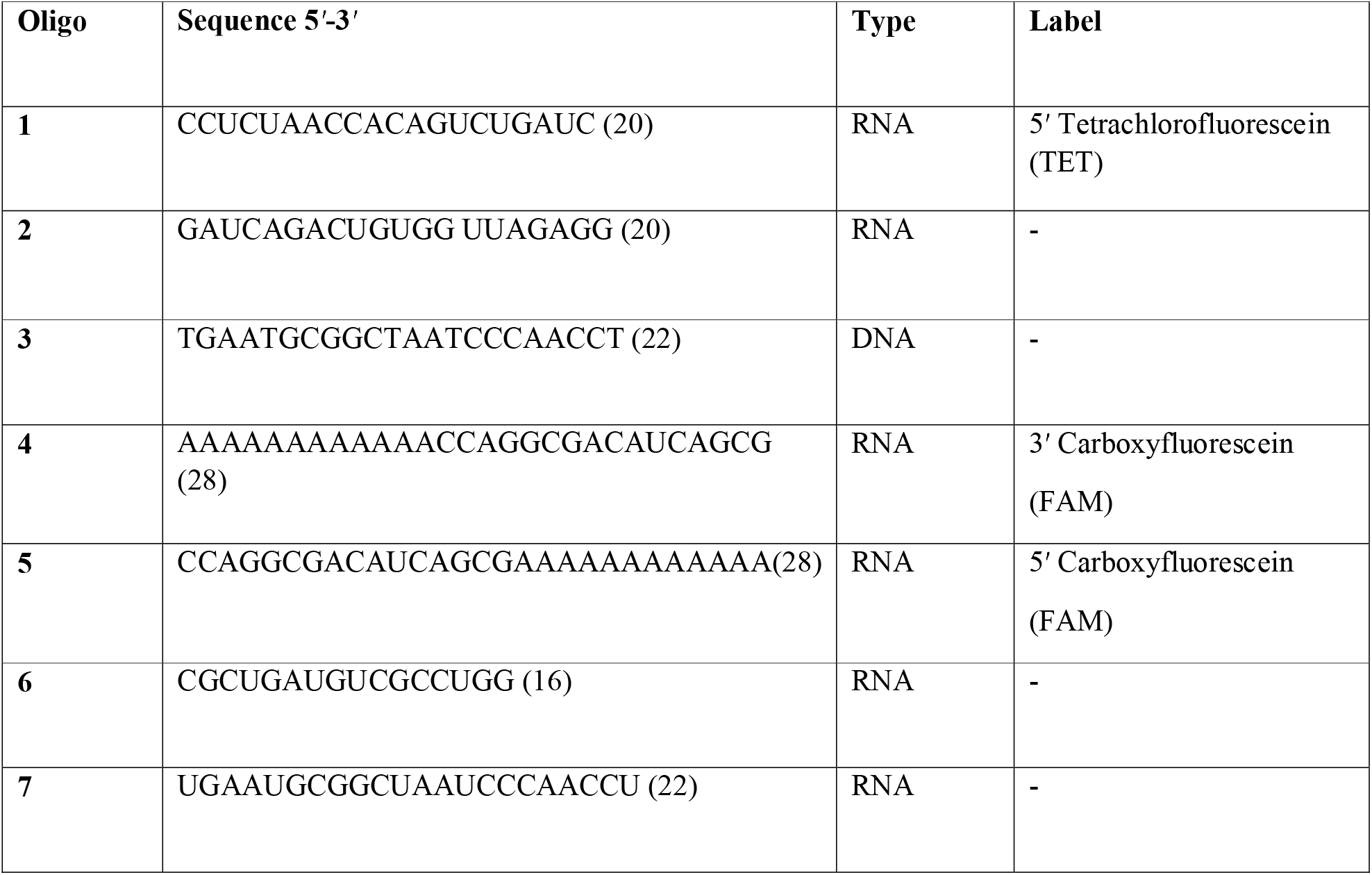
List of all the oligonucleotides used.

### Dynamic light scattering

100 µl of reaction mixture was prepared containing 0.8 μM or 3 μM 2C, 1 mg/ml SUVs, and TEV protease was incubated for 30 min at room temperature. Then, this mixture was transferred into a cuvette and the particle size was determined by dynamic light scattering using a Ζetasizer Nano-ZS (Malvern Instruments, Worcestershire, UK) at room temperature. To check the effect of ssDNA and ssRNA on clusteringssDNA oligo (5′ TGAATGCGGCTAATCCCAACCT 3′) (Eurofins Genomics) or ssRNA oligo (5′UGAAUGCGG CUAAUCCCAACCU 3′) (Eurofins Genomics) was added into the mix containing TEV protease, 1 mg/ml SUVs followed by addition of protein and the particle size was determined as described above.

### Sample preparation for cryo-electron tomography

A reaction mixture of 100 µl was prepared containing 3 μM His_6_MBP-2C, 1 mg/ml SUVs and 4 μM TEV protease was incubated for 30 min at room temperature. 3 μl of this mixture was blotted on a glow-discharged Quantifoil R 2/1 Copper 300 mesh grid (Electron Microscopy Sciences) and plunge-frozen in liquid ethane using a Vitrobot plunge freezer (ThermoFisher Scientific) at 22°C, 80% humidity, with blot force -20, blot time 2 seconds and wait time 40 seconds.

### Cryo-electron tomography data collection and processing

Tomographic tilt series were collected using a Titan Krios (ThermoFisher Scientific) operated at 300 kV, with a 100 μm objective aperture, Selectris energy filter and Falcon 4i detector (ThermoFisher Scientific). Tilt series were acquired in dose-symmetric mode, using SerialEM version 4. The following parameters were used for acquisition: 26,000 x nominal magnification corresponding to a specimen pixel size of 4.7 Å; defocus range of -3 μm to -8 µm, ± 63° tilt-range with tilt increment of 3°; total electron dose ∼116 e^−^/Å^2^. At each tilt angle, the exposure was saved as an EER movie. The EER files were converted to TIFF files using IMOD [81], gain-reference corrected, and motion corrected using MotionCor2 [82]. The tilt series were then processed and aligned using IMOD[81]. CTF was estimated using CTFFIND4 [83] and corrected using phase flipping as implemented in IMOD’s CTFPHASEFLIP.

The isotropic resolution and signal-to noise ratio of the tomograms were improved using the de-noising deep-learning software IsoNet [84]. The filtered tomograms were used for segmentation using Amira (Thermo Fisher Scientific) for the representation of SUVs and 2C. The average volume of 2C densities were subtracted from the segmentations using the surface area volume analysis available on Amira, thereafter the molecular weight was estimated assuming 825 Da/nm^3^.

### Colorimetric ATPase assay

Malachite Green Phosphate Assay Kit (MAK307, Sigma-Aldrich) was used for the colorimetric ATPase assay. For the enzymatic reaction 1.2 µM of His_6_MBP-2C (1-329), 2.9 µM TEV protease, 0.5 mg/ml SUV (POPC (75 mol %), POPE (25 mol %) and 0.4 mM ATP was mixed in buffer containing 40 mM HEPES pH7.5, 2 mM DTT and 12 mM NaCl and was incubated for 2 hours at 37 °C. The samples were diluted 20 times and plated in a 96 well plate and mixed with 20 µl working reagent supplied with the kit. The mixture was incubated at 37°C for 30 min after which the absorbance was measured at 620 nm according to the manufacturer’s instructions. A standard curve was prepared using the provided standard phosphate in the same buffer as the reaction. This standard curve was used to determine the phosphate liberated in the ATPase assay.

### Helicase and nuclease assays with RNA substrate

The single-stranded RNA (ssRNA) oligos (Eurofins Genomics) were annealed in molar ratio 1:1.5 (28 nucleotide labeled oligomer (5′ AAAAAAAAAAAACCAGGCGACAUCAGCG-FAM3′): 16 oligomer un-labeled (5′CGCUGAUGUCGCCUGG3 ′) in a buffer containing 10 mM HEPES-NaOH (pH 7.2) and 10 mM KCl. This mix was heated to 95°C for 1 minute and then cooled down to 22°C at the rate of 1°C per minute. The sample was then kept at 22°C for one hour. The thus prepared dsRNA with 5′ overhang was kept at 4°C overnight before freezing at -20°C. The dsRNA with 3′ overhang was made similarly using 28 nucleotide labeled oligomer (FAM-5′CCAGGCGACAUCAGCGAAAAAAAAAAAA3 ′) and mixing with 16 oligomer un-labeled (5′CGCUGAUGUCGCCUGG3′). Before starting the unwinding assay, 0.8 µM or 1.6 µM of MBP-2C was incubated with 0.8 mg/ml of SUVs (POPC (25 mol %), POPE (25 mol %) and 250 nM of TEV protease in reaction buffer containing 40 mM HEPES pH7.5, 2 mM DTT and 15mM NaCl, 0.5 µl RNAse inhibitor murine (NEB). This mixture was incubated for 90 min at 37°C. Then, to this mixture 20 nM dsRNA template, 400 nM ssRNA trap strand (28 nucleotide un-labeled oligomer). at 37°C for 20 min, followed by addition of 3.5 mM magnesium-ATP to start the reaction. For experiments comparing the effect of Mg^2+^ on the ribonuclease activity of 2C, 3.5 mM Mg^2+^ was added to the reaction. For experiments comparing the effect of 2C on ssRNA, 20 nM of each of 5′ labeled or 3′ labeled ssRNA was used. All the reaction were incubated for 2 hours before adding 5 µl of quenching buffer consisting of 100 mM Tris pH 7.5, 0.1% bromophenol blue, 1% SDS, 50 mM EDTA, and 50% glycerol. The samples were run on 16% native polyacrylamide gels for 2 hours at 100 V and scanned using an Amersham Typhoon fluorescence gel scanner. The gel scans were quantified using ImageJ, measuring the intensity of the bands at the ssRNA position as defined by the ssRNA control and subtracting the intensity of the dsRNA control as background.

### Luciferase assay

The P2-P3 region from poliovirus type 1 Mahoney (GenBank: V01149.1) was inserted into a vector containing *Renilla* luciferase reporter on the N-terminus. Truncated 2C versions (Δ2-11 and Δ2-40) and replication-defective 3D polymerase where the signature GDD [85] motif was changed to GAA were also constructed. The first glycine residue of 2C was not truncated in order to preserve the 3C protease cleavage site. The plasmids were linearized using ApaI restriction enzyme followed by *in vitro* transcription using MEGAscript™ T7 Transcription Kit (Invitrogen) according to the manufacturer’s protocol. The quality of the RNA was checked using 0.5% denaturing agarose gel. For transfection, 1 µg RNA was used and transfected in HeLa cells using TransIT®-mRNA Transfection Kit (mirusbio) according to the manufacturer’s instruction. The cells were harvested for luciferase assay after incubation at 37°C for 12 hrs. The same procedure was repeated for GFP-CVB3 replicon RNA which was used as transfection control (negative control for luciferase signal). The cells were lysed using lysis buffer supplied with Renilla Luciferase Assay Kit 2.0 (Biotium). 20 µl of cell lysate was mixed with 100 µl of freshly prepared working solution and the luminescence was recorded using a TECAN spectrophotometer.

### Negative-staining TEM

For negative-staining TEM, a reaction mixture of 100 µl was prepared containing 0.8 μM or 3 μM His_6_-MBP-2C, 1 mg/ml SUVs, and 4 μM TEV protease was incubated for 30 min at room temperature. From this mixture, 3 µl of sample was applied to a continuous carbon grid (Ted Pella, 300 mesh). The grid was then stained thrice for 10 s with 1.5% uranyl acetate drops, blotted and allowed to air dry. Images were taken at a Thermo Fisher Talos L120C operated at 120 kV, equipped with a 4,096 × 4,096-pixel Ceta camera. The nominal magnification was 36,000× pixel size.

### Inductively coupled plasma-optical emission spectrometry (ICP-OES)

To check the presence of zinc in our protein preparation, we used ICP-OES. 100 µM each of labeled and unlabeled His_6_-MBP-2C was mixed with 10 ml of 1% Suprapur® nitric acid (Merck Millipore). The mixture was filtered with a 0.2 µm filter. Then, the sample was loaded on the Agilent 5800 VDV ICP-OES (Agilent technologies, Santa Clara, California, US) machine and the emission corresponding to zinc was recorded (202 nm). Zinc acetate was used as a control.

### Mass Spectrometry of TEV protease

TEV protease was run on SDS-PAGE gel and the bands corresponding to the protease were cut from the gel and sent to the Proteomics Core Facility (University of Gothenburg) in 3% acetic acid for mass spectrometry and analysis of the peptides.

### Graphs and statistical procedure

Statistical significance tests were performed using student’s T-test as implemented in GraphPad Prism v10.0.1. Error bars represent standard deviation. P value style:; ns: p > 0.05; *: p < 0.05; **: p < 0.01; ***: p < 0.001; ****: p < 0.0001

## Data availability

The cryo-electron tomogram shown in Fig. 4C has been deposited to the Electron Microscopy Data Bank with accession code EMD-18615.

## Acknowledgements

We would like to thank Madeleine Ramstedt (Department of Chemistry, Umeå University) for the introduction to the Zetasizer Malvern Nanoz used for DLS measurements and Mikael Lindberg (Protein expertise platform, Umeå University, part of Protein Production Sweden) for cloning the poliovirus replicons. Electron microscopy was performed at the Umeå Center for Electron Microscopy (UCEM) a SciLifeLab National Cryo-EM facility part of the National Microscopy Infrastructure, NMI (VR-RFI 2016–00968). This project was funded by a Human Frontier Science Program Career Development Award (CDA00047/2017 C) to LAC, a Kempe foundations postdoctoral fellowship to MNS, an MSCA fellowship to HS (“MsInfection”, grant agreement ID: 101033469), the Knut and Alice Wallenberg Foundation (through the Wallenberg Center for Molecular Medicine Umeå), and the Swedish research council (grants 2018–05851 and 2021–01145).

---

**Figure S1:**
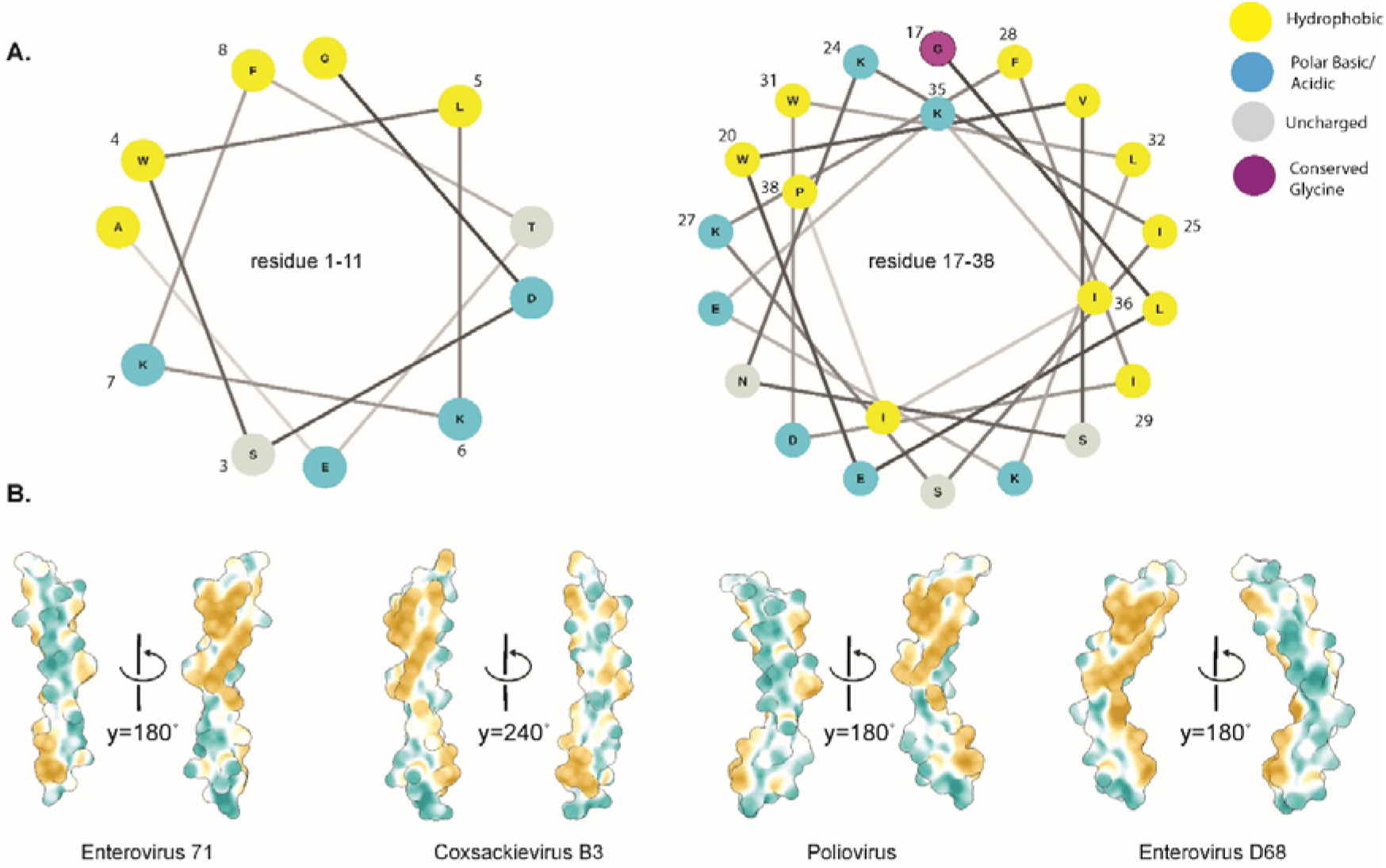
Amphipathic nature of the 2C membrane-binding domain. **(A)** Helical wheel of the AH1 and AH2 regions of poliovirus type 1 Mahoney 2C (GenBank: V01148.1) showing the distribution of hydrophobic and hydrophilic residues. Conserved residues are numbered. **(B)** Isosurface representations of Alphafold2-predicted structures of the MBD of different enterovirus genotypes showing the distribution of hydrophilic (blue) and hydrophobic (yellow) residues. For each structure prediction, two views related by the indicated rotation are shown.

**Figure S2:**
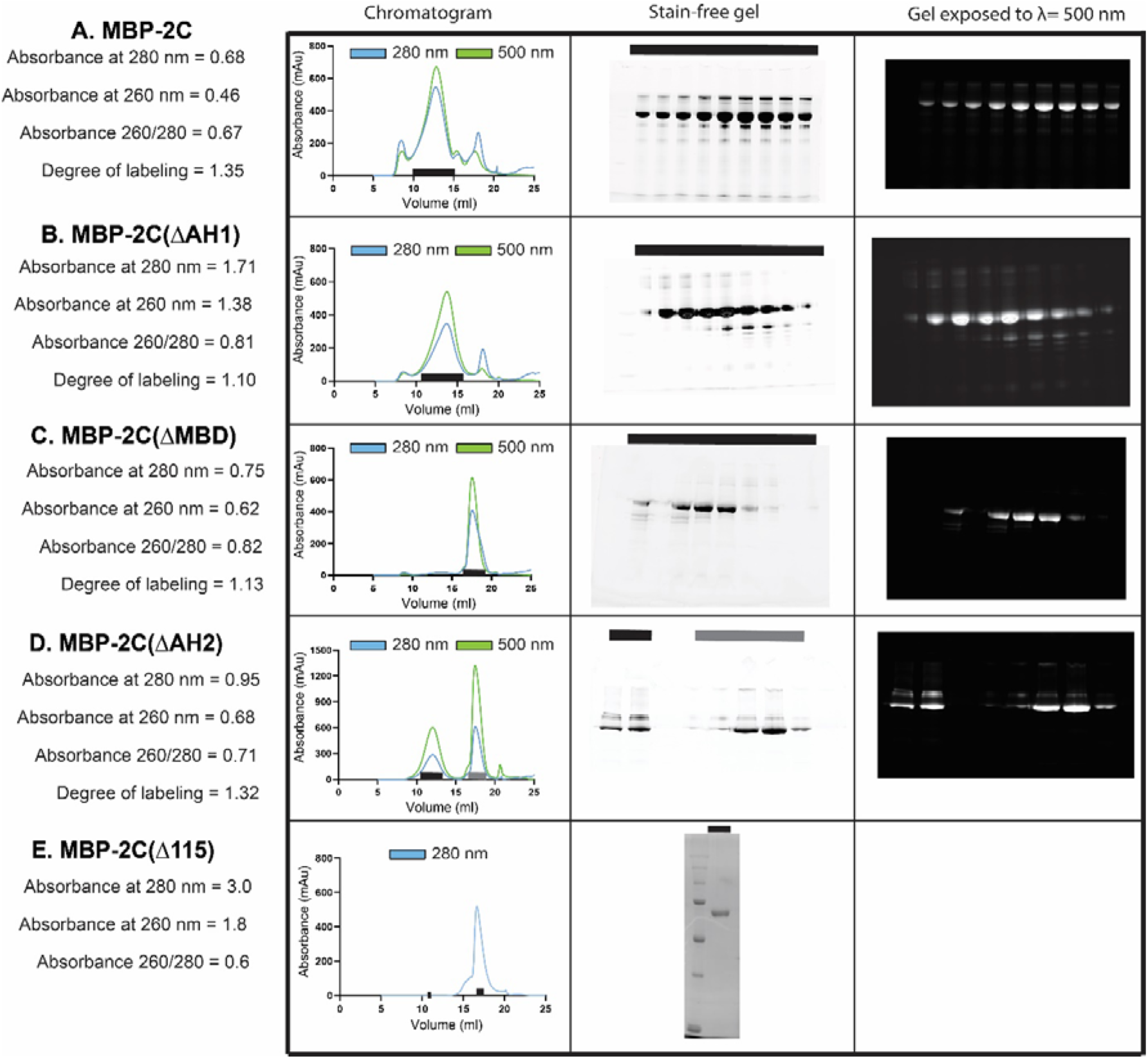
Degree of labeling, size-exclusion chromatography and SDS-PAGE gels for the fluorophore-labelled 2C constructs. **(A-D)** Absorbance at 280 nm, 260 nm, 260/280 ratio and degree of labeling for different constructs of MBP tagged 2C and the corresponding chromatogram obtained from Superose 6 Increase. The fractions that were run on an SDS-PAGE gel are marked on the chromatogram as solid lines. The SDS-PAGE gels were both exposed to UV for visualization of protein fractions and at = 500 nm to excite the ATTO 488 fluorophore to visualize the labeled protein. **(E)** MBP-2C(Δ115) Absorbance at 280 nm, 260 nm, and 260/280 ratio. The SEC chromatogram obtained from Superose 6 Increase and the corresponding stain free gel of the fraction marked on the chromatogram.

**Figure S3:**
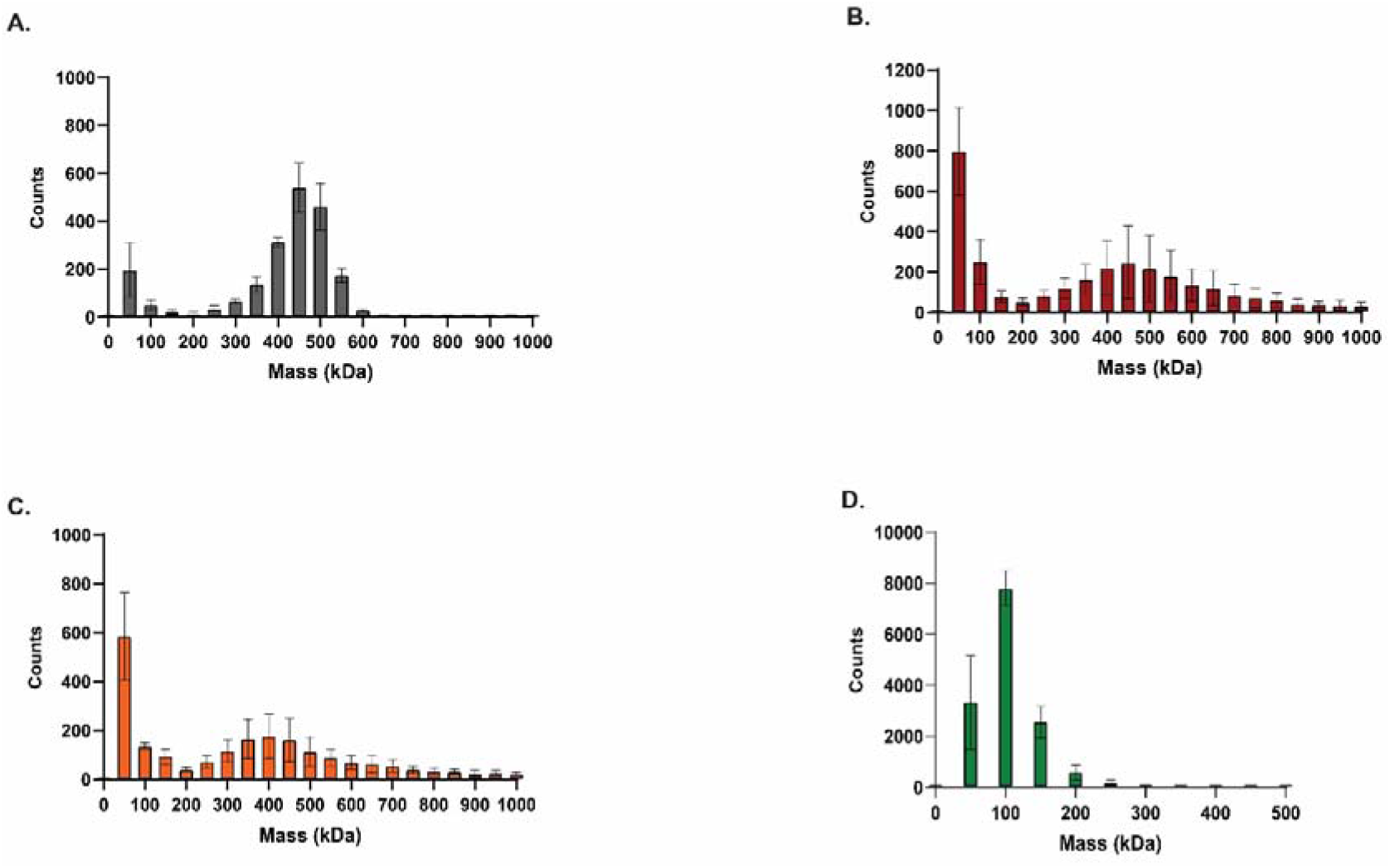
Mass photometry on apoferritin, MBP-2C, MBP-2C(ΔAH1) and MBP-2C(ΔAH2). Raw data (mean and standard deviation) from triplicates of mass photometry analysis of 100 nM each of **(A)** apoferritin, **(B)** MBP-2C, **(C)** MBP-(ΔAH1) and **(D)** MBP-(ΔAH2).

**Figure S4:**
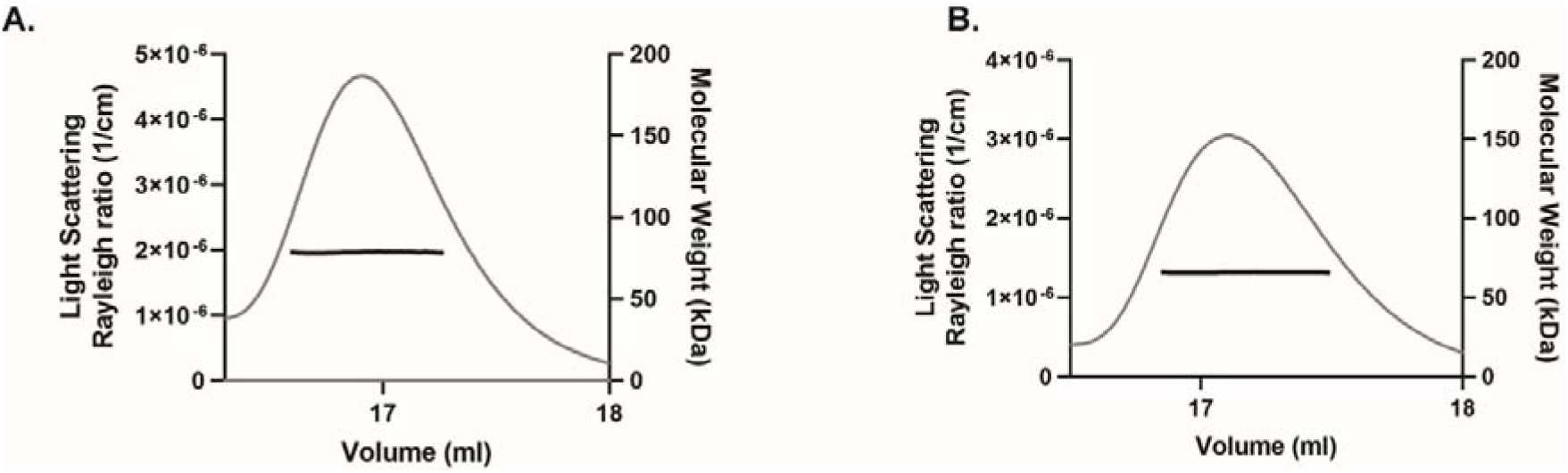
SEC-MALS mass analysis of MBP-fused, truncated 2C constructs. SEC-MALS data showing estimated molecular mass throughout the main elution peak of (**A**) MBP-2C(ΔMBD) and (**B**) MBP-2C(Δ115) at concentrations of 38 and 134μM, respectively. The calculated masses are shown as a thick black line. MBP-2C(ΔMBD) had an estimated mass of 78 kDa, whereas MBP-2C(Δ115) had an estimated mass of 66 kDa.

**Figure S5:**
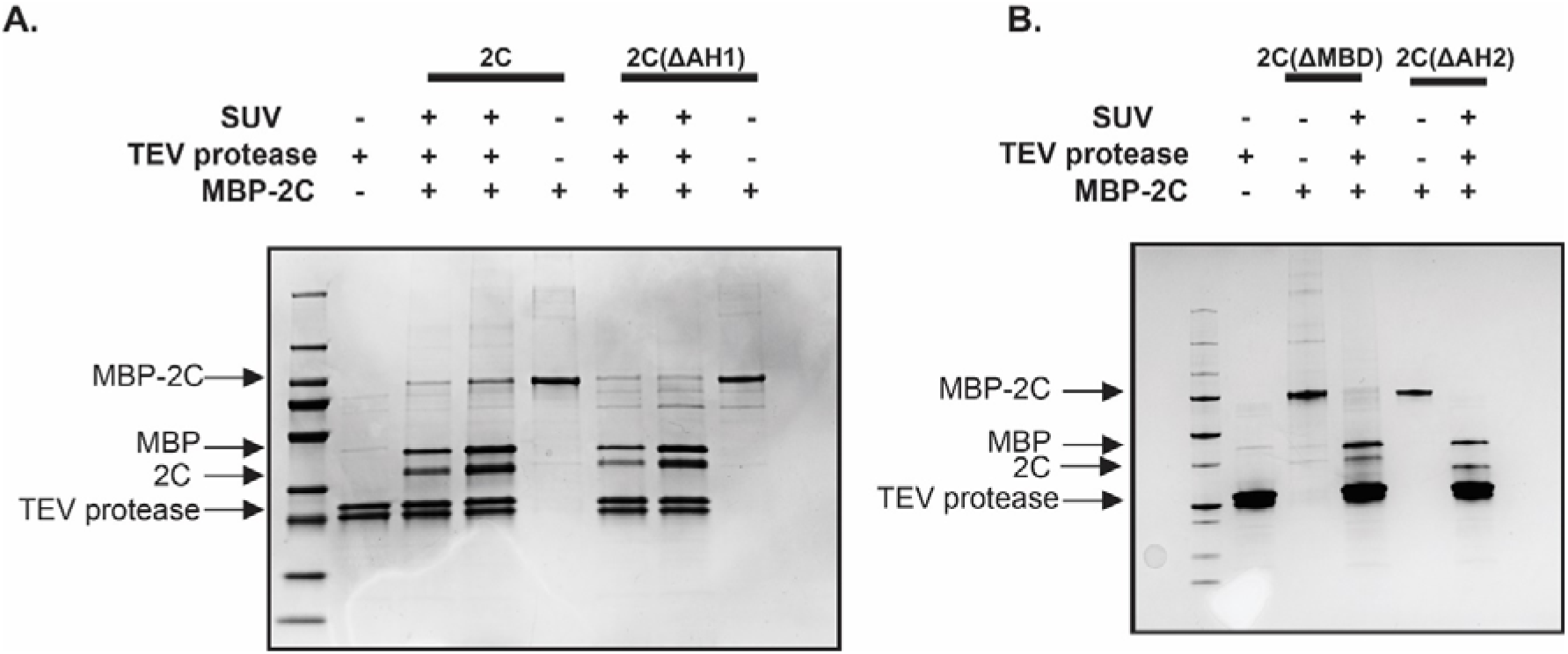
SDS-PAGE gel of material analyzed by flotation assay and DLS. **(A)** Lane 1: protein ladder. Lanes 2-8: Reactions including MBP-2C or MBP-2C(ΔAH1), and other components as indicated, at the end of the reaction. Lanes 3 and 6: 0.8 µM of the indicated 2C construct. Lanes 4-5 and 7-8: 3 µM of the indicated 2C construct. The two bands observed in the TEV protease prep were both confirmed to be TEV protease by mass spectrometry (Fig. S14). **(B)** As (A) but with 0.8 μM MBP-2C(ΔMBD) and MBP-2C(ΔAH2), as well as other components as indicated.

**Figure S6:**
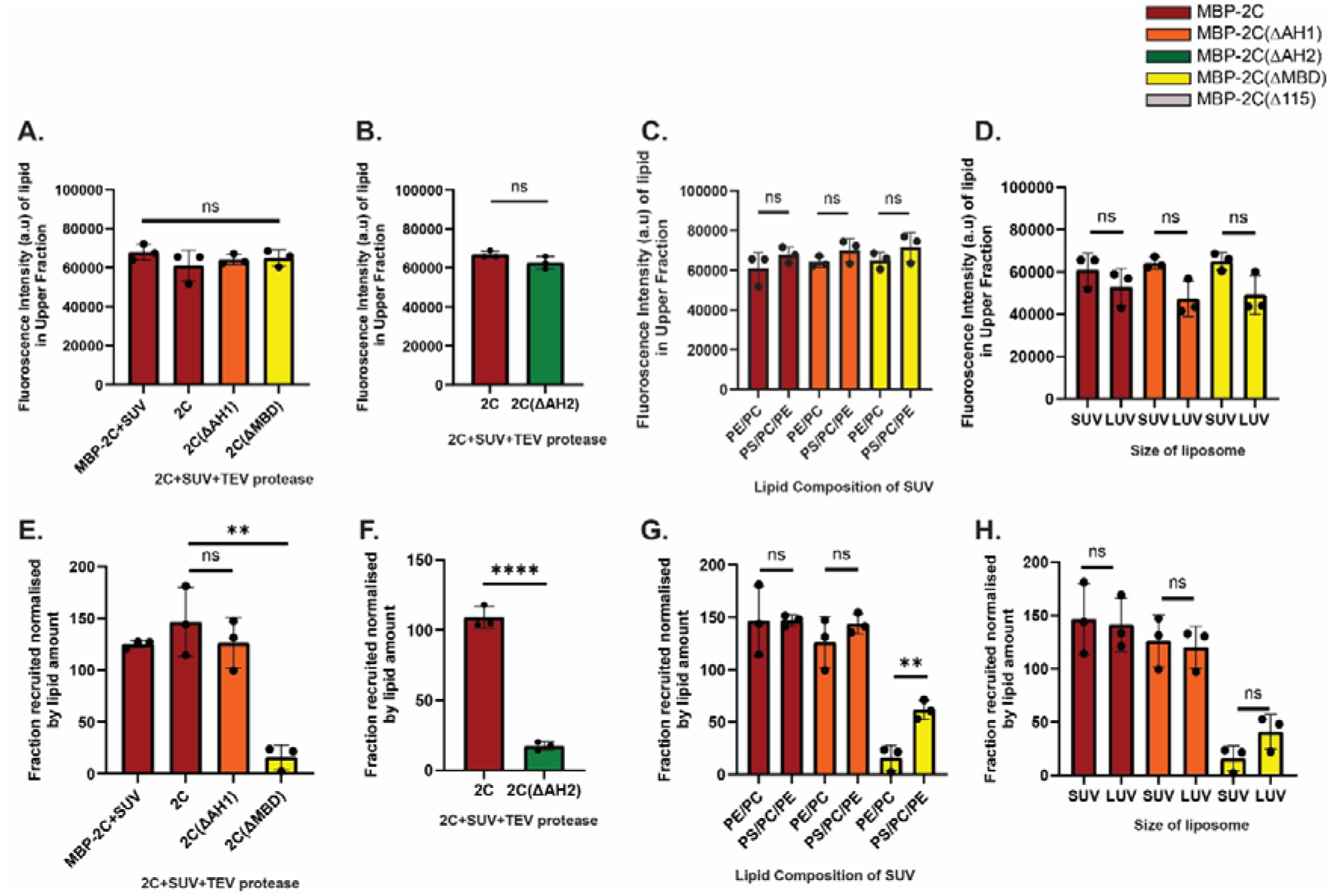
Quantitation of lipid fluorescence in flotation experiments. **(A-D)** Lipid fluorescence signal in the upper fraction of the assays shown in Fig. 3C-F. Error bars represent a standard deviation of three repeats of the experiment. Statistical significance by unpaired two-tailed Student’s t test; ns > 0.05. (**E-H**) Fraction 2C recruited to the upper fraction, normalized by amount of lipid in the upper fraction, for the assays shown in Fig. 3C-E. Error bars represent a standard deviation of three repeats of the experiment. Statistical significance by unpaired two-tailed Student’s t-test; ns: p>0.05, **: p<0.01.

**Figure S7:**
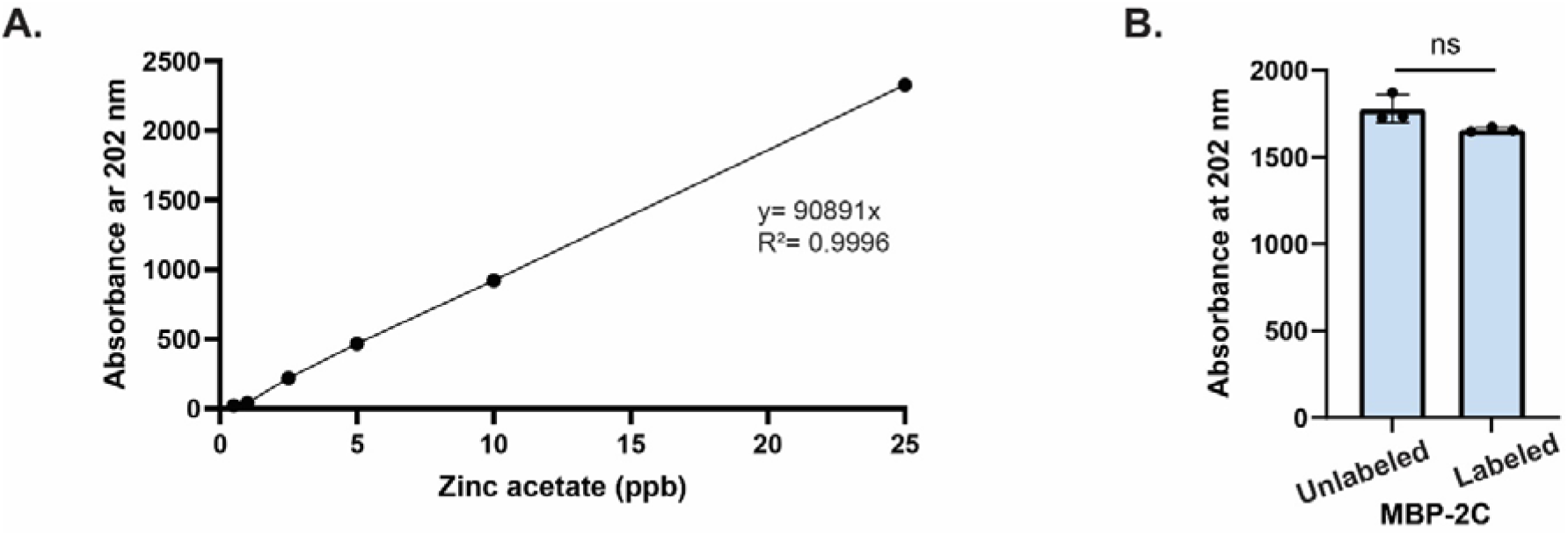
Zinc is detected in equal amount in fluorophore-labeled and unlabeled MBP-2C. **(A)** Standard curve with different concentrations of zinc acetate at 202 nm recorded using inductively coupled plasma-optical emission spectrometry (ICP-OES). **(B)** Emission at 202 nm corresponding to zinc, measured for labeled and unlabeled MBP-2C. Error bars represent a standard deviation of three repeats of the experiment. Statistical significance by unpaired two-tailed Student’s t-test; ns: p>0.05.

**Figure S8:**
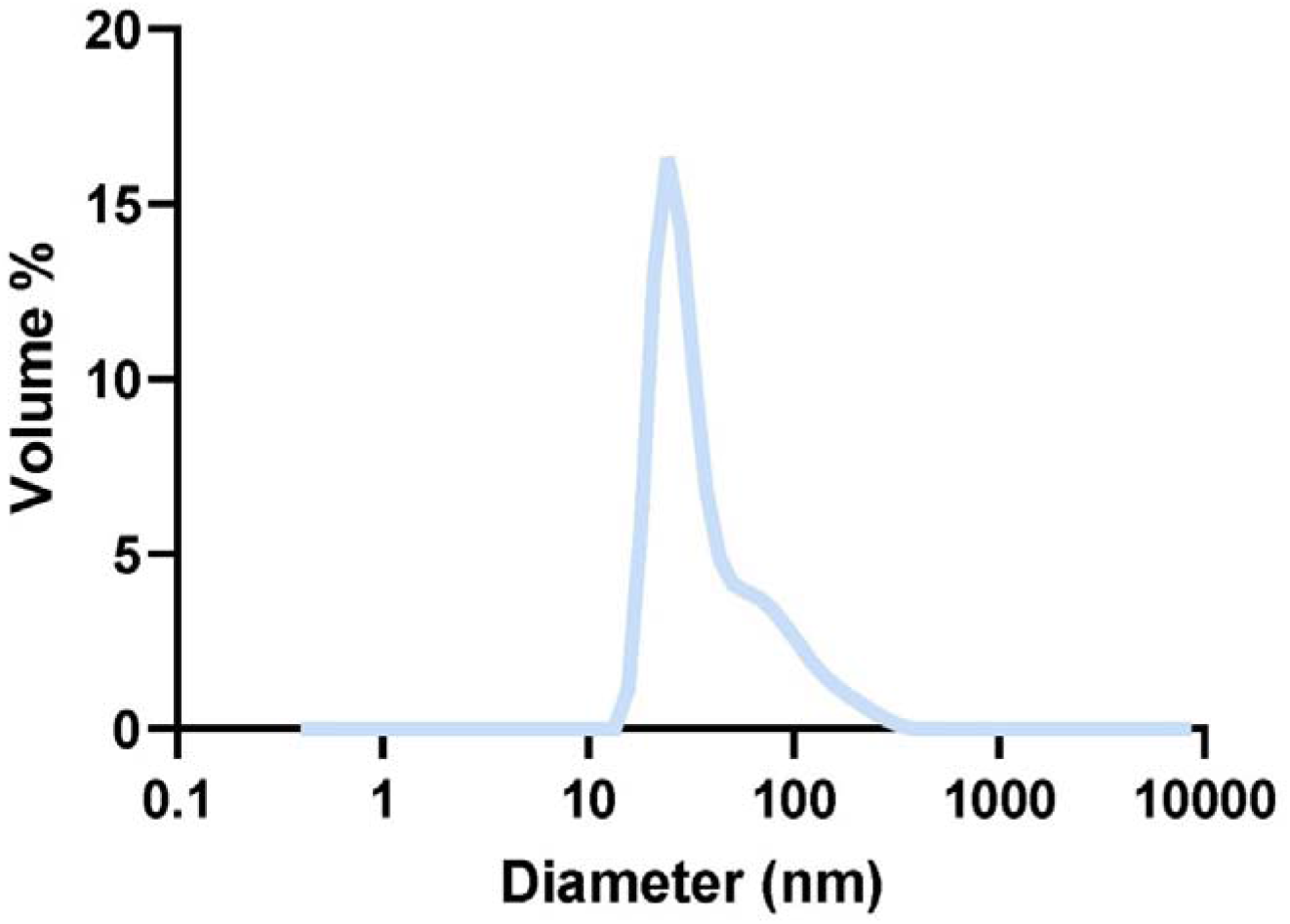
Size distribution of SUVs. Dynamic light scattering graph showing the size distribution of PE/PC SUVs (size distribution by volume). Average size of the SUVs was calculated to be 30 nm.

**Figure S9:**
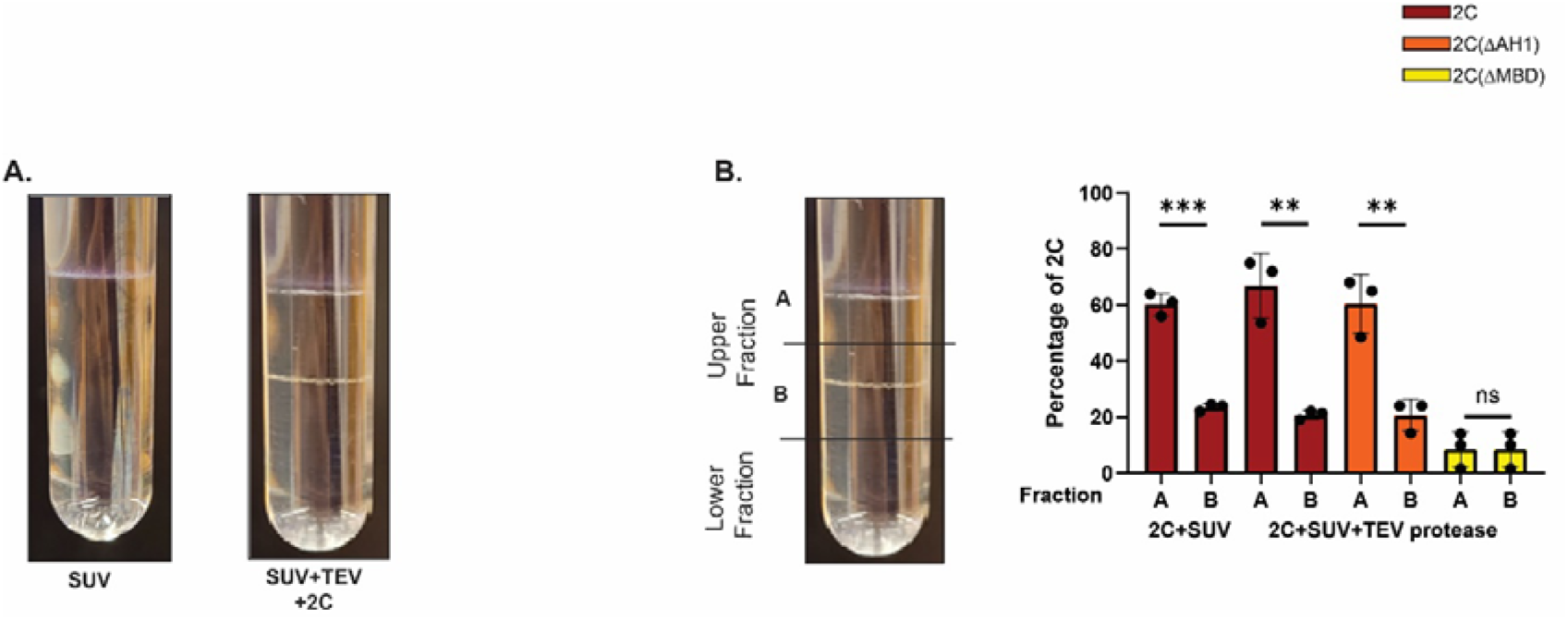
Two distinct lipid bands in the upper fraction in 2C flotation assays. **(A)** Photograph of representative centrifuge tubes after the flotation assay with SUVs only (single lipid band), and SUVs + TEV protease + MBP-2C (two lipid bands). **(B)** Photograph of centrifuge tube, indicating the upper fractions A, B, and the lower fraction. The percentage of the total 2C fluorescence in fractions A and B are shown. Error bars represent the standard deviation of three repeats of the experiment. Statistical significance by unpaired two-tailed Student’s t-test; **: p<0.01, ****: p<0.0001.

**Figure S10:**
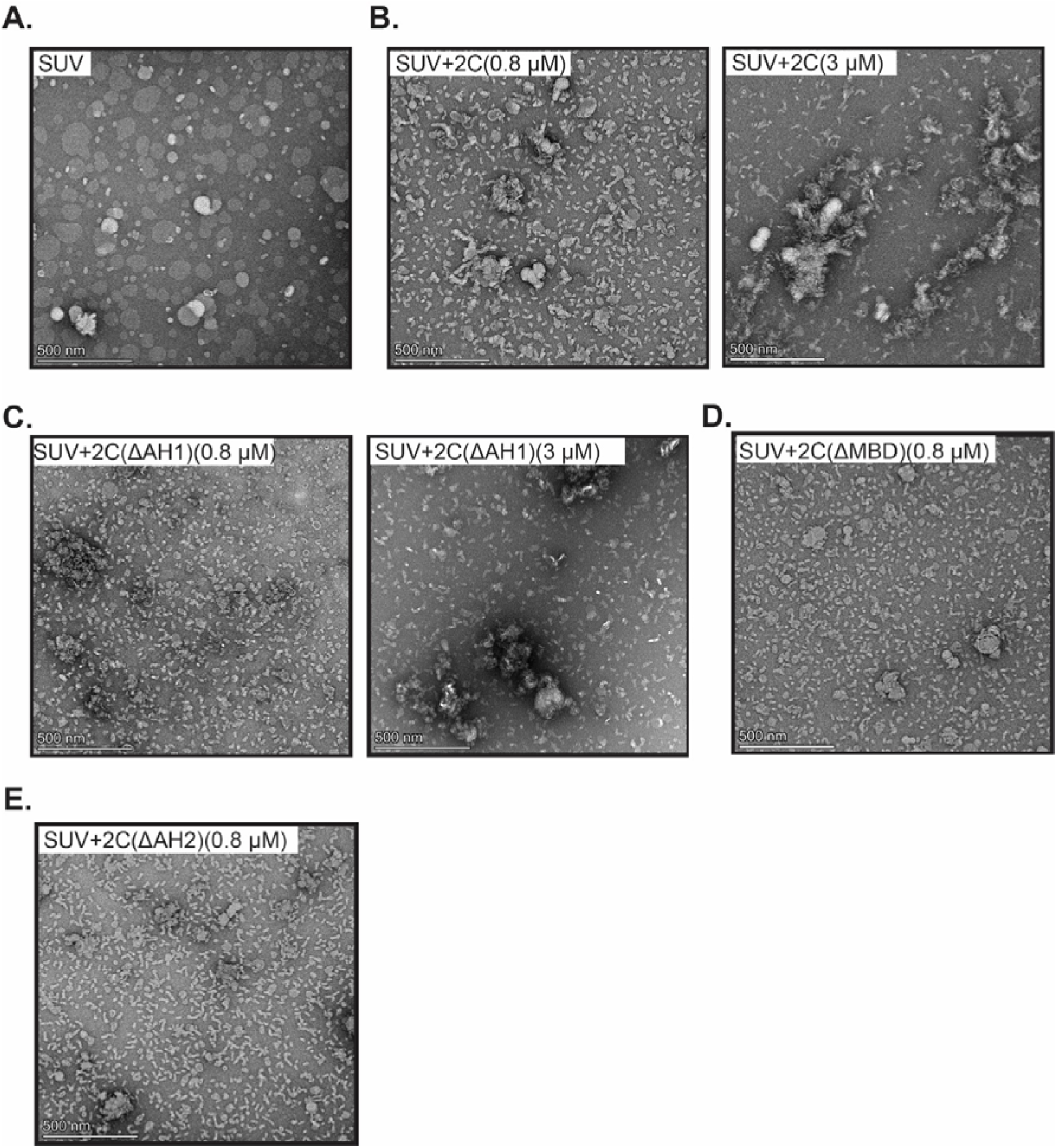
Negative staining electron microscopy of SUVs incubated with full-length and truncated 2C constructs. Representative images showing effect of different 2C constructs on SUVs **(A)** SUV only, **(B)** SUV+2C (0.8 µM and 3 µM), **(C)** SUV+2C(ΔAH1) (0.8 µM and 3 µM), **(D)** SUV+2C(ΔAH1) (0.8 µM), **(E)** SUV+2C(ΔAH2) (0.8 µM).

**Figure S11:**
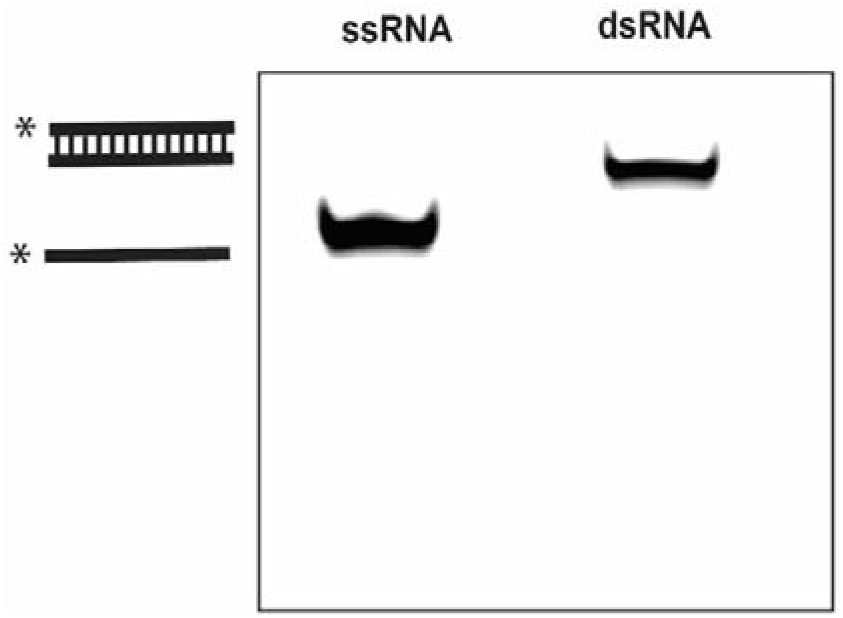
Characterization of fluorescent ssRNA and dsRNA. Fluorescence scan of non-denaturing polyacrylamide gel showing 5′TET-labelled ssRNA and 5′TET-labelled dsRNA.

**Figure S12:**
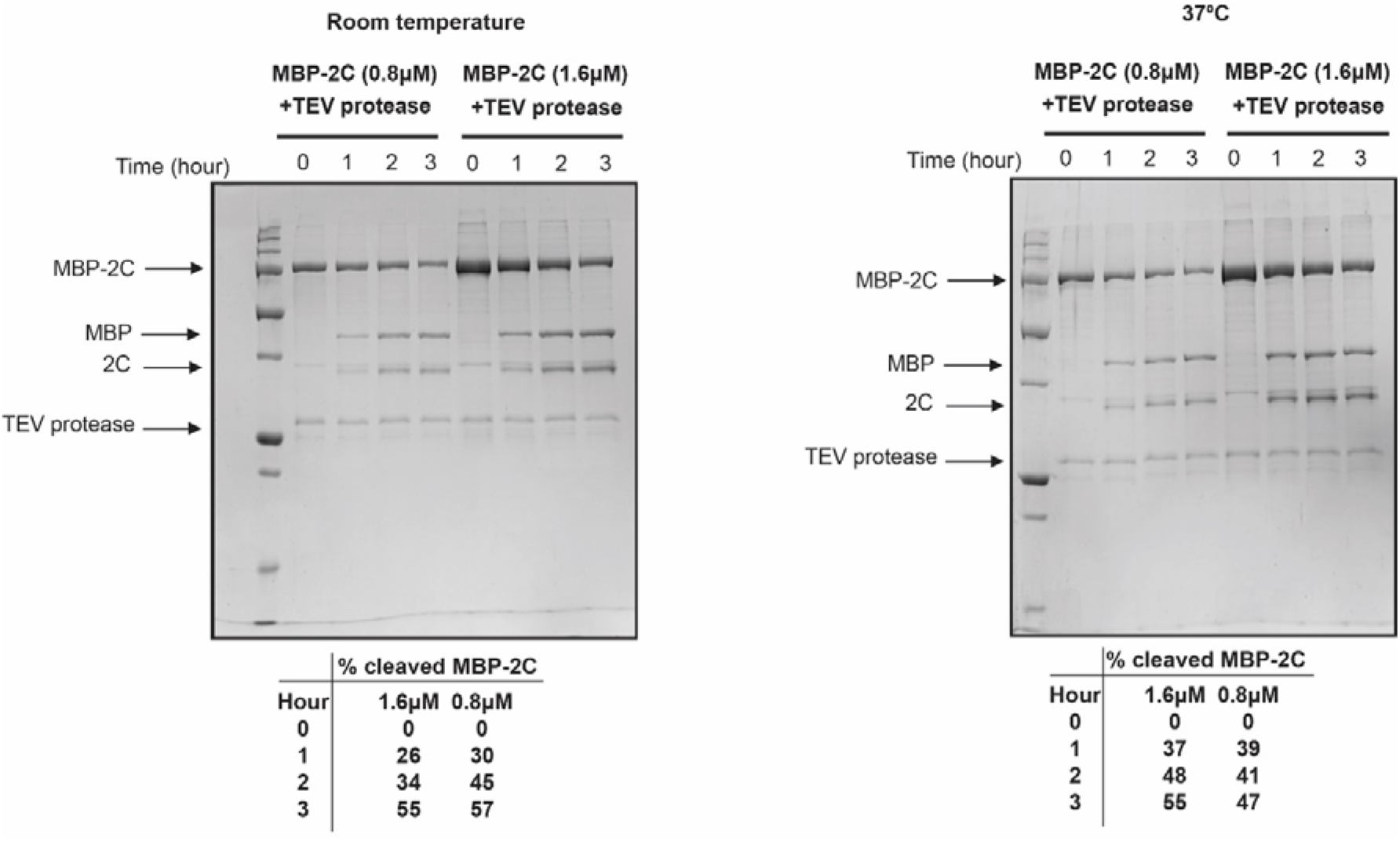
Time course of MBP-2C cleavage during helicase assay. SDS-PAGE gel showing cleavage of MBP-2C at different time points with 250 nM of TEV protease used in the dsRNA unwinding assay at room temperature and 37°C and the corresponding quantitation of the cleavage.

**Figure S13:**
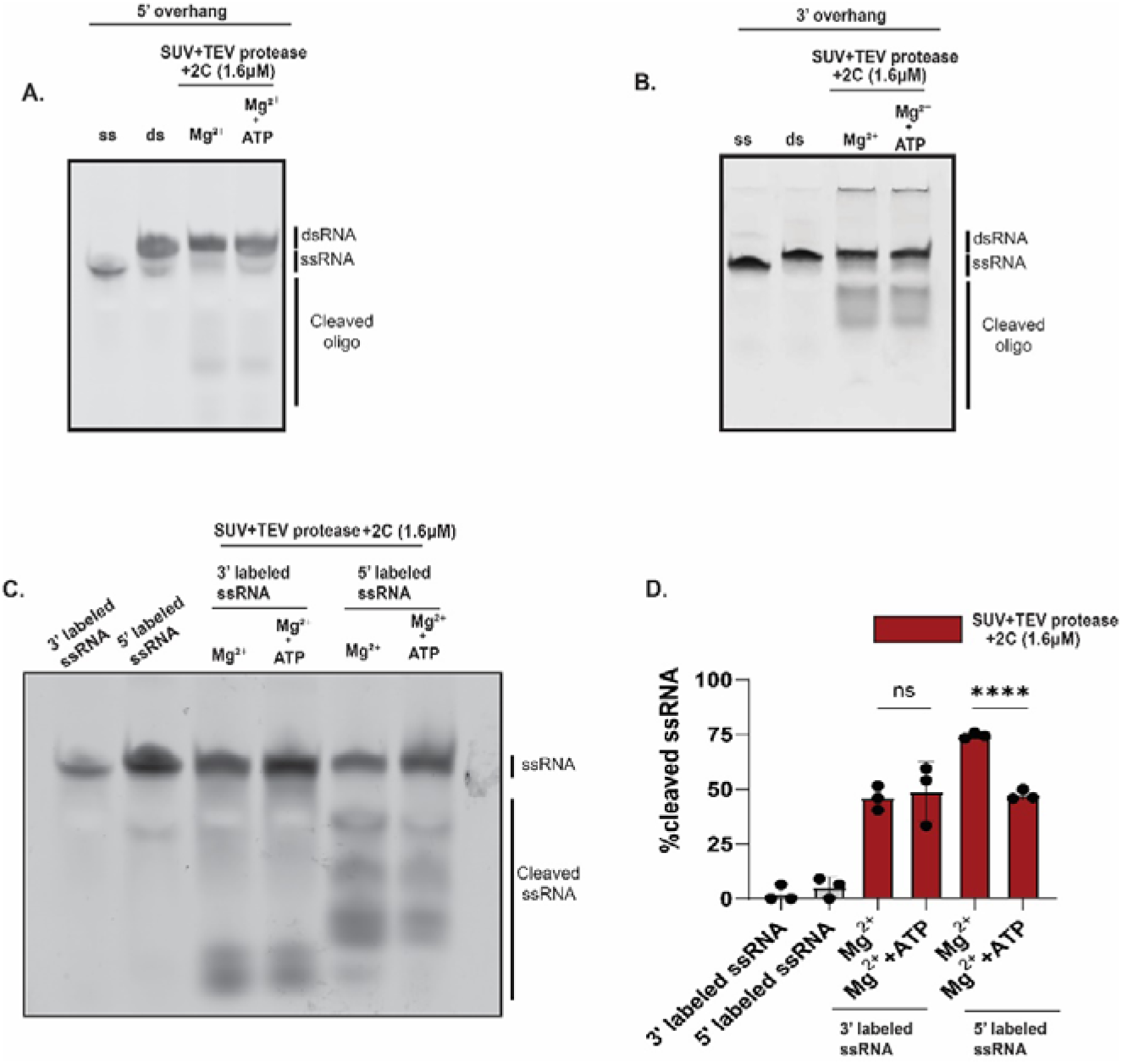
Ribonuclease activity of membrane-bound 2C is Mg^2+^-dependent but ATP-independent. **(A-B)** Fluorescence scans of native polyacrylamide gel for dsRNA substrate with 5′ (A) and 3′ (B) overhang. The first two lanes of each gel contain ssRNA control and dsRNA substrate. The following lanes correspond to the last lane in Fig. 6B,F, except that they have either Mg^2+^ or Mg^2+^+ATP added. Representative gels of four independent replicates are shown. **(C)** Experiment as (A) but using ssRNA substrates fluorophore-labeled at 3′ or 5′ end as indicated. A representative of three independent replicates is shown. **(D)** The percentage of total RNA signal present as cleaved oligo is plotted for lanes of the gel in (C), and two independent replicates of the same experiment. Dots represent individual experiments, bars the average and error bars one standard deviation. Statistical significance by unpaired two-tailed Student’s t-test. ns: p>0.05, ****: p<0.0001

**Figure S14:**
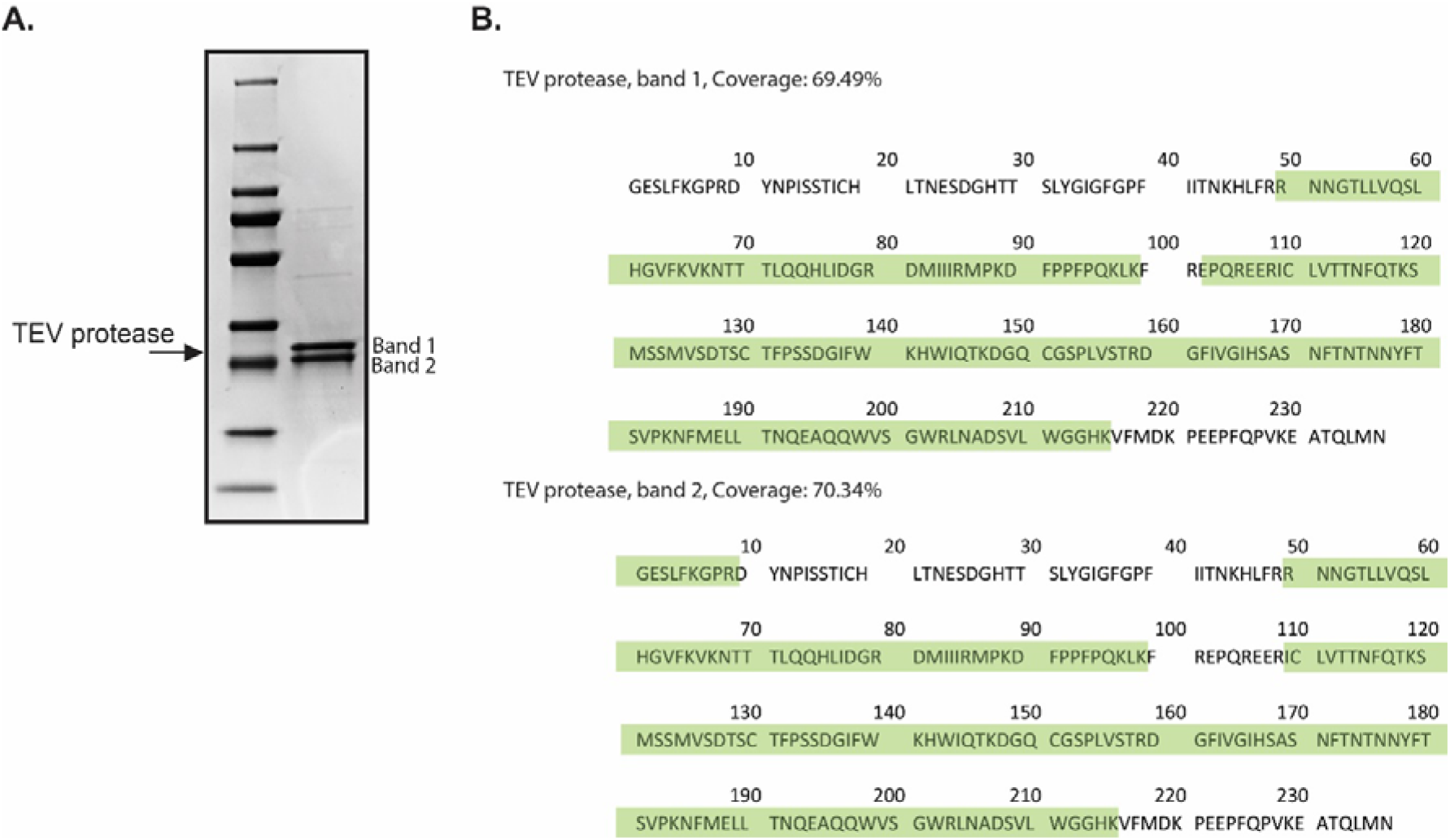
Mass spectrometry on two bands in the TEV protease prep. **(A)** SDS-PAGE gel showing 2 bands corresponding to TEV protease **(B)** Peptide coverage of TEV protease from in-gel digestion mass spectrometry of the excised bands from (A).

**Supplementary movie 1:** Slices through cryo-electron tomogram showing vesicle clustering by 2C. The movie loops consecutive xy-planes of the cryo-electron tomogram shown in Fig. 4C.

**Supplementary movie 2. Surface model of cryo-electron tomogram showing vesicle clustering by 2C.** The movie shows different views of the surface model of the tomogram shown in Fig. 4C.

